# Oxygen-dependent changes in HIF binding partners and post-translational modifications regulate stability and transcriptional activity

**DOI:** 10.1101/2020.11.12.379768

**Authors:** Leonard Daly, Philip J. Brownridge, Violaine Sée, Claire E. Eyers

## Abstract

Adaption of cells to low oxygen environments is an essential process mediated in part by the Hypoxia Inducible Factors (HIFs). Like other transcription factors, the stability and transcriptional activity of HIFs, and consequently the hypoxic response, are regulated by post-translational modification (PTM) and changes in biomolecular interactions. However, our current understanding of PTM-mediated regulation of HIFs is primarily based on *in vitro* protein fragment-based studies, with validation typically having been conducted by *in cellulo* fragment expression and hypoxia mimicking drugs. Consequently, we still lack an understanding of true oxygen deprivation signaling via HIFα. Using an immunoprecipitation-based, mass spectrometry approach, we characterize the regulation of *in cellulo* expressed full-length HIF-1α and HIF-2α, in terms of both PTM and binding partners, in response to normoxia (21% oxygen) and hypoxia (1% oxygen). These studies revealed that a change in oxygen tension significantly alters the complexity and composition of HIF-α protein interaction networks, with HIF-2α in particular having an extended hypoxia-induced interactome, most notably with mitochondrial-associated proteins. Both HIFα isoforms are heavily covalently modified: we define ~40 different sites of PTM on each of HIF-1α and HIF-2α, comprising 13 different PTM types, including multiple cysteine modifications and a highly unusual phosphocysteine. Over 80% of the PTMs identified are novel, and approximately half exhibit oxygen-dependency under these conditions. Combined with domain and evolutionary analysis of >225 vertebrate species, we validate Ser31 phosphorylation on HIF-1α as a regulator of transcription, and propose functional roles for Thr406, Thr528 and Ser581 on HIF-2α.

## INTRODUCTION

Oxygen is essential for all aerobic organisms to sustain cellular respiration and fuel vital enzymatic processes. Deprivation of the appropriate levels of oxygen *i.e.* hypoxia, can cause permanent damage to tissues and cells. Hence, organisms have developed strategies to sense, adapt and restore oxygen levels to maintain homoeostasis. Hypoxia-inducible factors (HIFs) are a family of transcription factors which function as central regulators of the hypoxic response (Wang *et al.* 1995 & Tian *et al.* 1997). Their discovery, and molecular investigation of their regulation and physiological roles, led to the Nobel prize in Physiology or Medicine being awarded in 2019 to G. Semenza, P. Ratcliffe and W. Kaelin.

Active HIF is a heterodimeric transcription factor, comprised of two subunits, an oxygen-sensitive α subunit, and a constitutively expressed β-subunit. The oxygen sensitivity of HIFα proteins is conferred by three enzymatic hydroxylation events, requiring oxygen as a co-factor. Under sufficient oxygen availability, two proline residues are hydroxylated (HIF-1α: Pro402/564, HIF-2α: Pro405/531), a process which triggers protein ubiquitination and proteasomal degradation (Ivan *et al.* 2001, Jaakkola *et al.* 2001 & Masson *et al.* 2001). Hydroxylation of an asparagine residue (HIF-1α: Asn803, HIF-2α: Asn847) also reduces the association of HIFα with p300/CBP and decreases transcriptional activity (Lando *et al.* 2002 & Lando *et al.* 2002). In hypoxia, HIFα hydroxylation is impaired and HIFα is thus no longer marked for degradation, allowing it to form a transcriptionally active heterodimer with HIF-1β. Although HIF-1α was the first HIFα isoform to be reported (Wang *et al.* 1995), the closely related HIF-2α (also termed Endothelial PAS domain protein-1 (EPAS1), HIF-1α Like Factor, HIF Related Factor, and Member of PAS Superfamily 2 (MOPS)) is known to play key and non-redundant roles during the cellular response to hypoxia (Tian *et al.* 1997). Whilst the two HIFα isoforms share ~50% sequence identity, there are notable differences in their roles and regulation, including their sensitivity to oxygen levels, sub-nuclear localization, target genes and promoter occupancy (Bracken *et al.* 2006, Mole *et al.* 2009, Taylor *et al.* 2016 & Smythies *et al.* 2019).

Many transcription factors are extensively regulated by dynamic, often reversible post-translational modifications (PTMs) (Lees-Miller *et al.* 1992, Noguchi *et al.* 1999, Lanucara *et al.* 2016 & Mylonis *et al.* 2006), with fine-tuning of their function being coordinated by multiple types of PTM. Similarly, for the HIF proteins, adaptation to hypoxia is controlled by the site and type of PTM, with different modifications influencing protein stability, localization, ability to form protein complexes, and to bind to DNA. To date, some 37 distinct covalent PTMs have been identified on HIF-1α, including phosphorylation (Mylonis *et al.* 2006), acetylation (Jeong *et al.* 2002), methylation (Kim *et al.* 2016), ubiquitination (Tanimoto *et al.* 2000), sumoylation (Bae *et al.* 2004) and nitrosylation (Li *et al.* 2007). However, these sites have typically been identified using fragment-based strategies, using recombinant HIFα fragments and/or modifying enzymes to identify PTMs *in vitro*. Moreover, the few studies that report cellular HIF-1α PTMs employ hypoxia-mimicking drugs (such as dimethyloxalylglycine (DMOG) and deferoxamine (DFO)) instead of oxygen-deprivation. Such drugs, while useful, are also known to regulate HIF hydroxylation patterns differently and influence a variety of other cellular signaling pathways (Tian *et al.* 2011, Ohyashiki *et al.* 2009 & Yu *et al.* 2011). Consequently, the majority of currently identified HIFα PTMs are biased, and confounded by lack of consideration towards full-length protein folding and regions of accessibility, and a lack of essential co-factor protein binding, a true understanding of how oxygen deprivation may affect global intracellular networks of cellular signaling. In addition, comprehensive comparative information on the PTM diversity and protein interaction networks of HIF-2α, either *in vitro* or in cells, is currently lacking.

Here, we report the first unbiased proteomics analysis to investigate the effect of oxygen deprivation (1% O_2_) in cells on their protein interaction networks and PTM status of HIF-1α and HIF-2α, with an emphasis on protein phosphorylation. To maximize protein coverage (and thus PTM site mapping), we exploited a complementary three protease strategy, achieving between 66% and 87% sequence coverage of the two HIFα isoforms. We identified a total of 41 and 39 PTMs for HIF-1α and HIF-2α respectively, with the vast majority (>80%) of these being novel, including phosphorylation, acetylation, methylation and cysteine-based modifications. Excitingly, we reveal a HIF-1α unique site of cysteine phosphorylation, pCys90. To evaluate the importance of these novel PTM sites, we investigated their evolutionarily conservation and explored functional effects of phosphosite amino acid substitutions. Using phospho-mimetic and phospho-null protein variants, we were able to define regulatory roles for some of these novel phosphorylation events, including Ser31 of HIF-1α and Thr406, Thr528 and Ser581 of HIF-2α, in HIFα stability and transcriptional activity.

## RESULTS

### Controlled expression of full-length HA-Clover-HIF-1α and HA-Clover-HIF-2α for immunoprecipitation and biochemical analysis

To obtain an unbiased analysis of all PTMs occurring on HIF-1α and HIF-2α subunits, as well as their binding partners, we devised a workflow for mass spectrometry (MS)-based proteomics analyses that would maximize HIF sequence coverage (Fig. 1A). One inherent problem with investigation of cellular HIFα proteins is the very low abundance of endogenous protein (estimated at ~11 ppm and ~6 ppm for HIF-1α and HIF-2α respectively, data from PAXdb (Wang *et al.* 2012)). Analysis is further complicated by the rapid (~5 min) turnover of HIFα under physiological (oxygenated) conditions (Moroz *et al.* 2009). To obtain enough material for MS-based PTM analysis and achieve efficient HIFα immunoprecipitation, we overexpressed HA-Clover tagged HIFα proteins (HA-Clover-HIFα) in HeLa cells and utilized GFP-Trap technology for protein enrichment (~70% efficiency; Supp. Fig. 1A). Transfection levels were optimized for maximal HIFα expression while minimizing apoptosis, and maintaining a degree of oxygen-dependent transcriptional activity (Fig 1B) as well as correct sub-nuclear localization (Fig 1C, Taylor *et al.* 2016). Final protein expression levels for IP were ~10 fold higher than the endogenous protein in 1% O_2_ (4 h incubation, Supp. Fig1. A/B). Despite the high expression levels, HA-Clover-HIFα proteins maintained a small but detectable difference in protein levels between normoxic and hypoxic conditions (~1.2 fold, measured by densitometry; Supp. Fig. 1A/B), in line with the small increase in transcriptional activity in hypoxia (Fig. 1B). Immunoblotting of the HA-Clover-HIF-2α eluant with the anti-HIF-2α antibody revealed two bands (Supp. Fig. 1A), initially suggesting homodimer formation between HA-Clover-HIF-2α and endogenous HIF-2α. However, the faster migrating band was also observed when identical membranes were probed with an anti-GFP antibody (Supp. Fig. 1C), suggesting that both bands derive from HA-Clover-HIF-2α. Although it is possible that the faster migrating band is a C-terminal cleavage product, the apparent mass (~130 kDa) equates to the theoretically calculated HA-Clover-HIF-2α molecular mass of ~125 kDa, suggesting instead that the slower migrating form of HIF-2α likely arises due to extensive PTM. Interestingly, the faster migrating (presumably hypo-modified) protein band for HA-Clover-HIF-1α was not observed at either O_2_ tensions in the eluent or when probing for GFP, indicating this assumed PTM-modified form(s) of HIF-1α occurs rapidly and independent of O_2_ (Supp. Fig. 1A/D).

**Figure 1:**
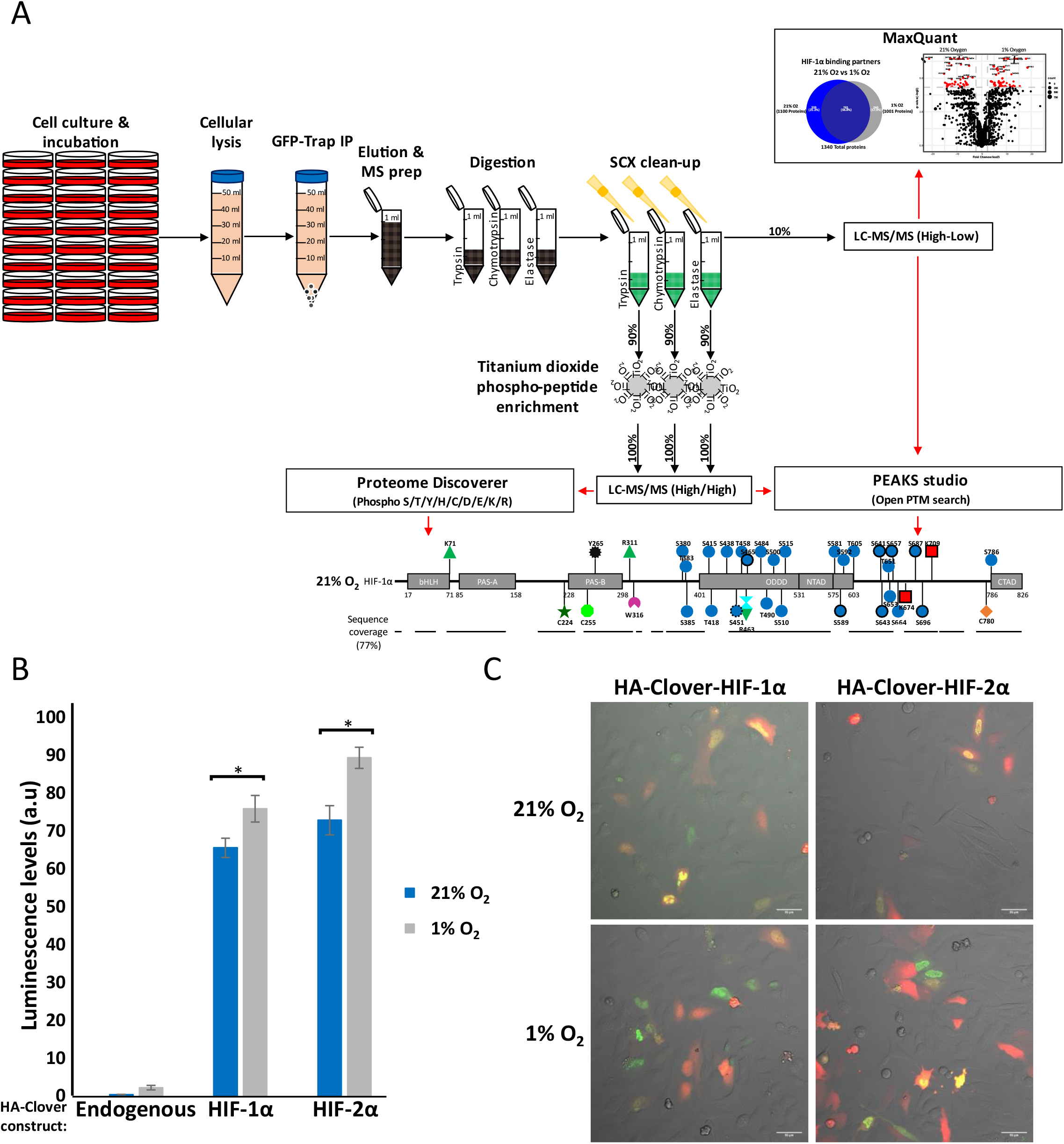
Optimised workflow for mass spectrometry analysis of HIFα from HeLa cells. A) Schematic representation of the workflow, which includes the main steps of immunoprecipitation (IP), clean-up/enrichment and data analysis strategy. B) Assessment of transcriptional activity upon HA-Clover-HIFα overexpression. An HRE-Luciferase reporter construct was co-transfected together with HIFa at high expression level (see methods, 1:1 HA-Clover-HIFα:HRE-Luciferase ratio) and endpoint luminometry readings were taken post 4 h incubation at 21% or 1% O_2_, n=3, *=*p*-value<0.05. C) Assessment of nuclear localization with high HA-Clover-HIFα expression. HeLa cells were transfected at high expression level (1:1 ratio HA-Clover-HIFα:pcDNA-mRUBY) and confocal images taken 24 h after transfection and upon 4 h incubation at 21% or 1% O_2_. A scale bar of 50 μm is shown. Representative of n=3.

### Hypoxia promotes isoform-specific HIF-α interaction networks

Perhaps not unexpectedly, the complement of proteins interacting with either HIF-1α (Fig 2) or HIF-2α (Fig 3) were significantly altered by hypoxia. Filtering for interactors that were identified in both biological replicates (Supp. Table. 1), we identified 579 and 726 interactors for HIF-1α and HIF-2α respectively (equating to ~45-50% of the total interactome for each isoform) that were specific to a given O_2_ tension (Fig. 2A and Fig. 3A). Interestingly, hypoxia had a much greater influence on the interactome of HIF-2α than HIF-1α (Fig. 2A versus Fig. 3A), with a significantly greater proportion of the HIF-2α interaction network being identified only in 1% O_2_. Although 240 proteins were identified as hypoxia-specific HIF-1α binding partners (24% of the total number of binding proteins in 1% O_2_, 18% of the total HIF-1α interactome), we identified 642 proteins as hypoxic-specific binding partners of HIF-2α, equating to 45% of the total proteins identified under these conditions (42% of the total HIF-2α interactome), suggesting a greater sensitivity of HIF-2α to oxygen tension-mediated protein complex formation. Interrogation of oxygen specific interactions of the two HIF isoforms reveals only part of the biological story; numerous proteins were identified irrespective of O_2_ tension (overlap region Fig. 2A and Fig. 3A; HIF-1α 761 (56.8% of total interactome) and HIF-2α 781 (51.8% of total interactome)), but at levels that varied as a consequence of O_2_ tension. We therefore evaluated the relative fold change in levels of protein binding partners as a function of O_2_ tension, using label-free quantification (LFQ) (Fig. 2B, Fig. 3B; Supp. Table. 2). Of the proteins identified in both normoxia and hypoxia, HIF-1α had 49 (~6%), and HIF-2α had 195 (~25%) interactors that were present at significantly different levels when cells were exposed to different O_2_ tensions.

**Figure 2:**
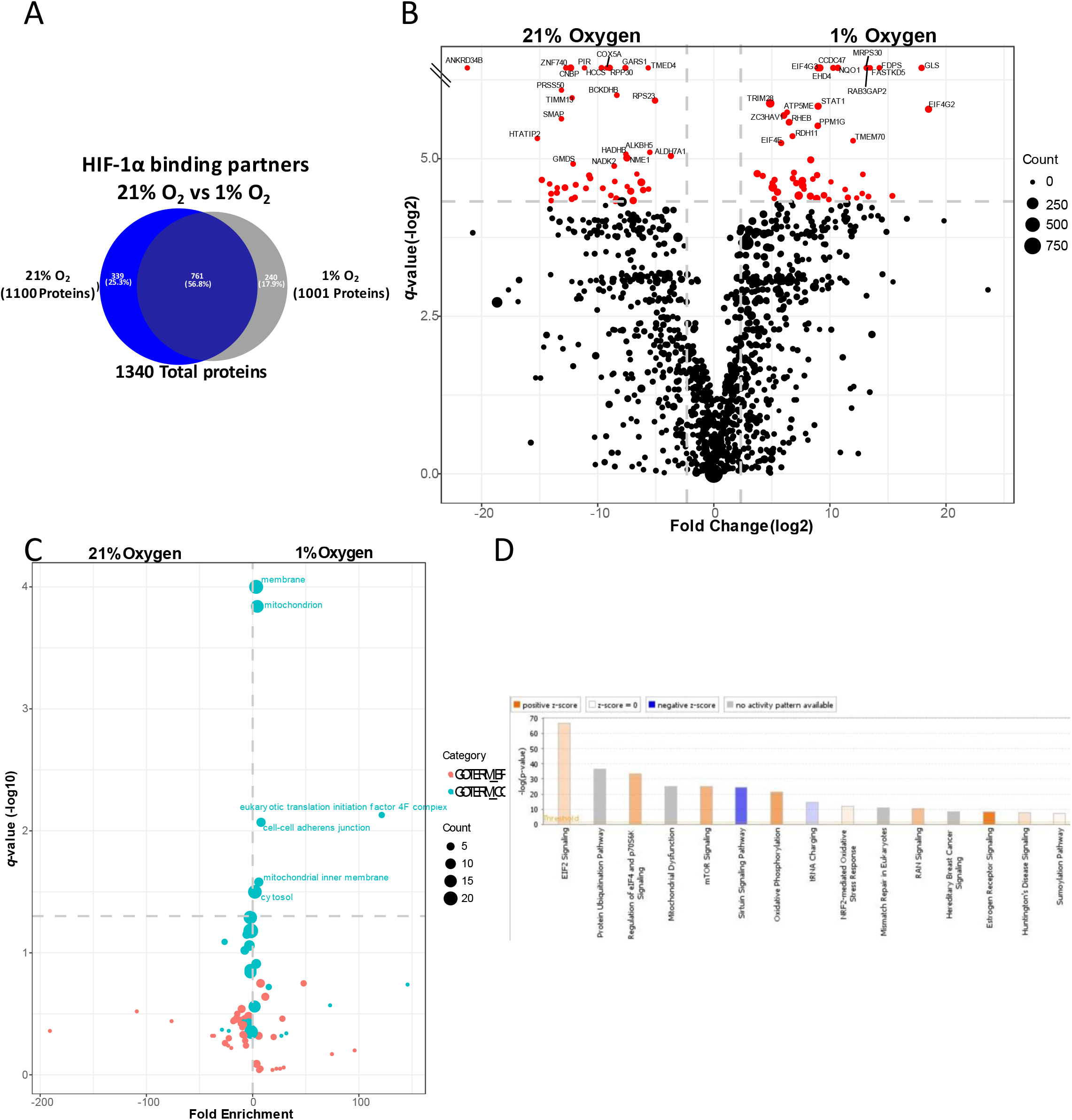
Hypoxia-mediated regulation of the HA-Clover-HIF-1α interactome. A) Overlap of all protein interactors (number and %) identified at 21% vs 1% O_2_. Proteins were maintained in the dataset if they were observed in both biological replicates (1% FDR cutoff following background subtraction). Blue = 21% O_2_; Grey = 1% O_2_, n=2. B) Volcano plot depicting results of the label free protein quantification, shown as –log2(*q*-value) vs log2(1%/21% abundance fold change). Replicate data was internally normalized respective to HA-Clover-HIF1α abundance. LHS (negative values) more abundant in 21% O_2_, RHS (positive values) more abundant in 1% O_2_. Dot size equates to the combined number of MS/MS events for a protein across all replicates. Black = *q*-value>0.05, Red = *q*-value<0.05, Top 20 *q*-value identifications for both O_2_ conditions are labelled with their gene name. Grey dashed lines are at 5-fold change and *q*-value = 0.05. *p*-value correction = FDR permutation based, 250 replicates, n=2. C) GO term enrichment analysis using DAVID of interactors significantly regulated by O_2_ tension (*q*-value <0.05). BP = Biological process; CC = cellular compartment. Dot size is equivalent to the number of proteins within a category. LHS (negative values) more abundant in 21% O_2_, RHS (positive values) more abundant in 1% O_2_. Top 20 *q*-value identifications for both O_2_ conditions are labelled. *p*-value correction = Benjamini-Hochberg, n=2. D) Oxygen-dependency, interactor pathway analysis. LFQ data was averaged between replicates and a 1%/21% abundance ratio determined for each protein and analyzed using IPA software. The top 15 most significantly identified pathways are displayed (−log_10_p-value), bars are colored dependent on whether they are predicted to be activated (orange) or inactivated (blue) by 1% O_2_. White = no predicted difference in activation, grey = does not have prediction data associated. A threshold line at p=0.05 is displayed, n=2.

**Figure 3:**
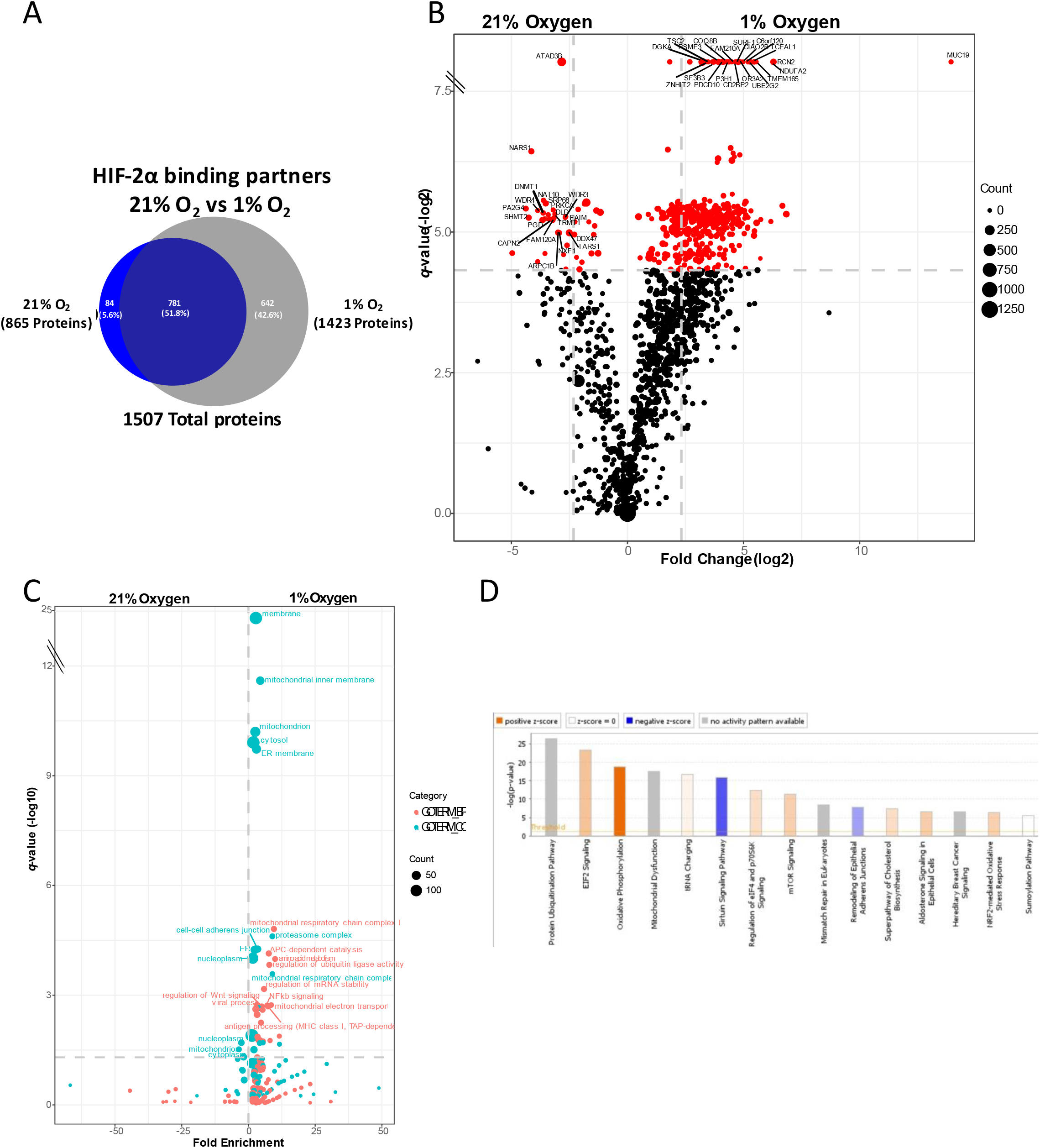
Hypoxia-mediated regulation of the HA-Clover-HIF-2α interactome. A) Overlap of all protein interactors (number and %) identified at 21% vs 1% O_2_. Proteins were maintained in the dataset if they were observed in both biological replicates (1% FDR cutoff following background subtraction). Blue = 21% O_2_; Grey = 1% O_2_, n=2. B) Volcano plot depicting results of the label free protein quantification, shown as –log2(*q*-value) vs log2(1%/21% abundance fold change). Replicate data was internally normalized respective to HA-Clover-HIF2α abundance. LHS (negative values) more abundant in 21% O_2_, RHS (positive values) more abundant in 1% O_2_. Dot size equates to the combined number of MS/MS events for a protein across all replicates. Black = *q*-value>0.05, Red = *q*-value<0.05. Top 20 *q*-value identifications for both O_2_ conditions are labelled with their gene name. Grey dashed lines are at 5-fold change and *q*-value = 0.05. *p*-value correction = FDR permutation based, 250 replicates, n=2. C) GO term enrichment analysis using DAVID of interactors significantly regulated by O_2_ tension (*q*-value <0.05). BP = Biological process; CC = cellular compartment. Dot size is equivalent to the number of proteins within a category. LHS (negative values) more abundant in 21% O_2_, RHS (positive values) more abundant in 1% O_2_. Top 20 *q*-value identifications for both O_2_ conditions are labelled. *p*-value correction = Benjamini-Hochberg, n=2. D) Oxygen-dependency, interactor pathway analysis. LFQ data was averaged between replicates and a 1%/21% abundance ratio determined for each protein and analyzed by IPA software. The top 15 most significantly identified pathways are displayed (-log_10_*p*-value), bars are colored dependent on whether they are predicted to be activated (orange) or inactivated (blue) by 1% O_2_. White = no predicted difference in activation, grey = does not have prediction data associated. A threshold line at p=0.05 is displayed, n=2.

A total of 97 HIF-1α binding proteins were significantly regulated by O_2_ tension (*q*-value <0.05), 47 were more reduced or absent in hypoxia, while 50 were at lower levels or absent under normoxic conditions (Fig. 2B). Of these significantly regulated proteins, all exhibited at least a 10-fold increase in O_2_ tension-regulated abundance. Although we determined 350 proteins in total to be differentially bound with a fold change of >100 (168 and 182 more abundant in normoxia and hypoxia respectively), only 77 were statistically different (*q*-value <0.05). The likely explanation to large fold changes that lack significance is due to protein identification in 1 replicate and imputed values in the 2^nd^ replicate (i.e. considered O_2_ tension specific in Fig. 2A), given that an additional 457 proteins were considered O_2_ tension specific in Fig. 2A yet had a *q*-value > 0.05; resulting in large variance.

For HIF-2α, 424 proteins were quantified as being of significantly different abundance at the two O_2_ tensions, with only 43 of these being elevated in normoxia (Fig. 3B). Of the 381 proteins significantly elevated in 1% O_2_, 315 exhibited a greater than 5-fold increase in abundance. The single largest oxygen-dependent change in HIF-2α interactors were tyrosine-protein phosphatase non-receptor type 3 (PTPN3, P26045), which was elevated ~31 fold under normoxic conditions, while mucin-19 (MUC19, Q7Z5P9) increased over 15,000 fold in hypoxia (the fold change of which is hard to define in absolute terms, given the imputation of values and the likelihood that mucin-19 is a true hypoxia dependent binding partner). The significant reduction in binding of HIF-2α to PTPN3 (directly or indirectly) in 1% O_2_ may suggest important roles in hypoxia-mediated regulation of phosphorylation status across this network, although we did not identify any sites of Tyr phosphorylation on HIF-2α.

Gene Ontology (GO) analysis of the identified significantly O_2_-dependent HIF interacting proteins (Fig. 2C & Fig. 3C) revealed enrichment of mitochondrial proteins (GO:0005739) for both HIF-1α and HIF-2α, with 26 within this category [21% O_2_: 10 proteins (~21% of significant proteins), *q*-value = 0.09; 1% O_2_: 16 proteins (~32% of significant proteins), *q*-value = 1.44e-4] for HIF-1α, and 81 [21% O_2_: 11 proteins (~26% of significant proteins), *q*-value = 0.03; 1% O_2_: 70 proteins (~18% of significant proteins), *q*-value = 6.87e-11] for HIF-2α (Supp. Table 3). Interestingly, ~18% of all identified HIF-1α and HIF-2α binding partners (independent of oxygen tension) were associated with the mitochondria (271 for HIF-1α, *q*-value = 3.94e-48, 292 for HIF-2α, *q*-value = 9.35e-51,Supp. Table. 3), suggesting that at least a proportion of both HIFα isoforms is likely associated with this organelle regardless of O_2_ tension, supporting previous findings (Briston *et al.* 2011 & Concolino *et al.* 2018).

To better understand oxygen-dependent involvement of HIF-1α and HIF-2α in cellular signaling, we used Ingenuity Pathway Analysis (IPA) software to explore enrichment of isoform and O_2_ tension-specific binding partners in different signaling pathways (Fig. 2D, Fig. 3D, Supp. Table. 4). Protein ubiquitination and EIF2 signaling were the top two most enriched pathways for both HIF isoforms. Although components of the EIF2 pathway were common for HIF-1/2α, both being predicted to be activated under hypoxic conditions (z-scores of 1.033 and 2.137 respectively), many more proteins within the EIF2 signaling pathway were identified in the HIF-1α interactome (HIF-1α 108 proteins, HIF-2α 61 proteins). Many (12 of the top 15) pathways identified between HIF-1α and HIF-2α were identical, with most predicted to be activated by hypoxia. Major activated pathways include mTOR signaling (z-scores = 2.132 and 1.633), regulation of eIF4 and p70S6K signaling (z-scores = 2.000 and 1.155) and oxidative phosphorylation (z-scores = 2.469 and 5.191 for HIF-1α and HIF-2α respectively). Interestingly, the only pathway within the top 15 predicted to be inactivated by hypoxia for both HIFα isoforms is Sirtuin signaling, with z-scores of −2.143 and −1.808 for HIF-1α and HIF-2α respectively.

We additionally analyzed all significant pathways identified by IPA analysis in terms of the predicted (in)activation by hypoxia (Supp. Table. 4). Both HIF-1α and HIF-2α have PPAR (peroxisome proliferator-activated receptor) signaling as the most inactivated pathway upon hypoxia treatment (HIF-1α: −log_10_(*p*-value) = 4.45, z-score = −3.441; HIF-2α: −log_10_(*p*-value) = 2.84, z-score = −2.673), and is the only shared pathway between isoforms in the top 5 most inactivated pathways. While the single most activated pathway by hypoxia was different between HIF-1α and HIF-2α, 3 of the top 5 pathways were identical: IL-8 (interleukin-8) signaling (HIF-1α: −log_10_(*p*-value) = 3.26, z-score = 4.082; HIF-2α: −log_10_(*p*-value) = 2.31, z-score = 3.13), Gα 12/13 signaling (HIF-1α: −log_10_(*p*-value) = 1.37, z-score = 3.742; HIF-2α: −log_10_(*p*-value) = 1.59, z-score = 3.207), and Estrogen receptor signaling (HIF-1α: −log_10_(*p*-value) = 8.27, z-score = 3.539; HIF-2α: −log_10_(*p*-value) = 5.08, z-score = 3.244). Overall, these data suggest that the major O_2_-sensitive pathways that regulate, or are regulated by, the two HIFα isoform are similar.

### HIFα isoforms have highly distinct interaction networks for a given oxygen tension

Given the distinct regulation and response to hypoxia of the two HIFα isoforms, particularly considering their relatively high sequence homology, we next compared HIF-1α and HIF-2α protein networks across the two different O_2_ tensions, evaluating those proteins that were specific to normoxia, hypoxia, or O_2_-independent (Fig. 4, Supp. Table. 5).

**Figure 4:**
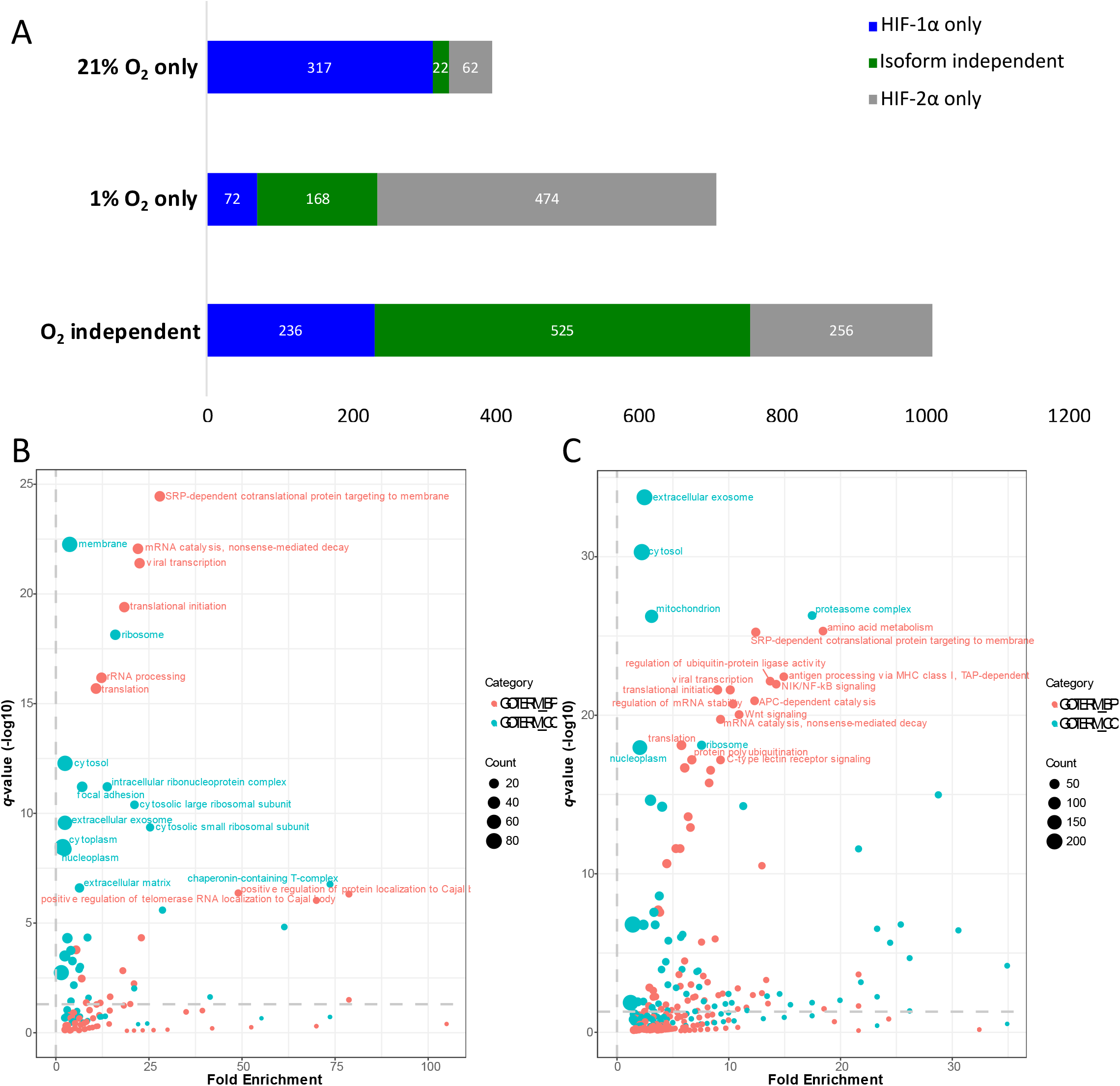
The interactomes of HIF-1α and HIF-2α are distinct for the same O_2_ tensions. A) Comparison of HIFα interactomes at the same O_2_ tension. Exact numbers of interactors under each of the different conditions are included in each section. Blue = HIF-1α specific interactors; Grey = HIF-2α specific interactors; Green = protein binding partners of both HIF-1/2α. Data filtered as in Fig 2A/3A, n=2. B) GO-term enrichment analysis of proteins identified as binding partners for both HIF-1α and HIF-2α under 1% O_2_ conditions (green section of (A), 168 proteins), filtered as in Fig 2C/3C. Dot size is proportional to number of proteins identified within that GO-term. Color indicates the respective GO-term, orange = BP (biological process), blue = CC (cellular compartment). Top 20 *q*-value identifications for both GO-terms are labelled. *p*-value correction = Benjamini-Hochberg, n=2. C) Go-term enrichment analysis of proteins identified as O_2_-independent interactors of both HIF-1α and HIF-2α (525 proteins, displayed as in Fig 4B), n=2.

Considering all the protein binding partners identified for both isoforms that were specific for 21% O_2_ (21% O_2_ only), only 5% (22 out of 401 total proteins identified) were common between the HIFα isoforms (Fig 4A). No significant enrichment in terms of biological process, or cellular compartment was identified for these 22 proteins (Supp. Table 6), suggesting that their regulation under normoxic conditions is isoform specific. In contrast, a greater proportion (24%; 168 out of 714) of the hypoxic-specific protein interactors were common for both isoforms. GO analysis of these 168 proteins revealed significant enrichment in proteins associated with telomere regulation: positive regulation of telomerase RNA localization to Cajal body (GO:1904874), *q*-value = 4.25e-7; positive regulation of protein localization to Cajal body (GO:1904871), *q*-value = 4.75e-7; positive regulation of establishment of protein localization to telomere (GO:1904851), *q*-value = 9.43e-7; nuclear chromosome telomeric region (GO:0000784), *q*-value =4.62e-5; positive regulation of telomere maintenance via telomerase (GO:0032212), *q*-value = 4.71e-5 (Fig. 4B; Supp. Table. 6). Identification of translation (GO:0006412, *q*-value = 2.06e-16), translation initiation (GO:0006413, *q*-value = 3.97e-20), ribosome (GO:0005840, *q*-value = 7.20e-19) and both small (GO:0022627)/large (GO:0022625) cytosolic ribosomal subunits (*q*-values = 4.41e-10 and 4.04e-11 respectively) as enriched biological processes also suggests a common role for both of these isoforms in hypoxic-mediated regulation of translational control. This is in line with our finding that proteins involved in EIF2 signaling are activated in hypoxic conditions in the interactomes of both HIF isoforms (Fig. 2D/3D) and with previous work from others (Uniacke *et al.* 2012). Interestingly, among the binding partners identified as O_2_ independent, 52% (525 out of 1016) were common between both HIFα isoforms (but not observed in negative control experiments). Although binding partners with roles in translation were identified for both isoforms irrespective of O_2_ tension, proteasome complex (GO:0000502) was one of the most significantly regulated terms (Fig. 4C, *q*-value = 5.08e-27), highlighting the well described and essential role of the proteasome in HIFα regulation. In agreement with our previous findings, mitochondrion (GO:0005739) was the 5^th^ most significantly enriched term for proteins identified as O_2_ independent binding partners, with 118 proteins matching this GO term (~23% of all proteins, *q-*value = 5.73e-27). Regulation of amino acid metabolism (*q-*value = 4.94e-26) also featured in this dataset, as did a number of cell signaling processes.

To further explore these, IPA was again used to investigate the potential role of these two HIF isoforms in different signaling pathways in an O_2_-dependent manner (Supp. Table. 7). Using hierarchal clustering of significantly enriched pathways, we found that the signaling pathways identified by HIFα isoforms were more similar for the same O_2_ tension than for the same isoform in different O_2_ tensions, suggesting O_2_ tension is a more significant regulator of cell signaling than isoform specificity (Supp. Fig. 2). As demonstrated above (Fig 2D/3D), EIF2 signaling and protein ubiquitination pathways were enriched for both isoforms, irrespective of O_2_ condition. IPA analysis indicated potential hypoxia-mediated crosstalk with RAN signaling for both HIFα isoforms (Fig. 2D/3D, Supp. Table. 7), possibly via differential nuclear shuttling during hypoxia. RAN GTPase is known to be required for nuclear export of the telomerase protein TERT (Haendeler *et al.* 2003 & Seimiya *et al.* 2000), a process which is regulated during oxygen stress. It is plausible therefore, based on our findings, that HIF-1/2α play roles in mediating these processes. Additionally, Wnt (*q-*value = 9.13e-21), MAPK (*q*-value = 2.29e-11), NF-κB (*q-*value = 1.10e-22), mTOR (−log(*p*-value) = 16.90 and 9.77 respectively) and NRF2 mediated oxidative stress response (−log(*p*-value) = 9.72 and 7.48 respectively) signaling pathways were all identified as major oxygen independent pathways for which both HIF-α isoforms have known binding partners (Figs. 4B/C and Supp. Table 6/7), supporting previously identified roles for these pathways in regulating cellular HIF (Xu *et al.* 2017, Sang *et al.* 2003, D’Ignazio *et al.* 2016, Cam *et al.* 2010 & Kim *et al.* 2011).

### HIF-1α and HIF-2α proteins are extensively modified in both normoxia and hypoxia

Like other transcription factors HIF proteins are known to be regulated by PTMs. However, the majority of studies to date seeking to understand PTM-mediated regulation of HIF proteins have employed *in vitro* assays, often using protein sub-domains. Thus, to understand the extent of cellular HIF-1α and HIF-2α PTMs in the context of O_2_ availability, we undertook comprehensive PTM mapping of immunoprecipitated HIF-1/2α proteins at different O_2_ tensions, using three complementary proteases to maximize HIF sequence coverage (Supp. Table 8). All PTMs discovered were cross-referenced against the UniProt (Consortium 2018) and PhosphositePlus (Hornbeck *et al.* 2012) databases, to ascertain novelty (Supp. Table. 9, Fig. 5). With a view to investigate potential fundamental and evolved regulatory mechanisms, the PTM sites identified were additionally evaluated for evolutionary conservation against 250 HIF-1α and 225 HIF-2α non-redundant vertebrate species (Supp. Table. 9, Supp. File. 1/2).

**Figure 5:**
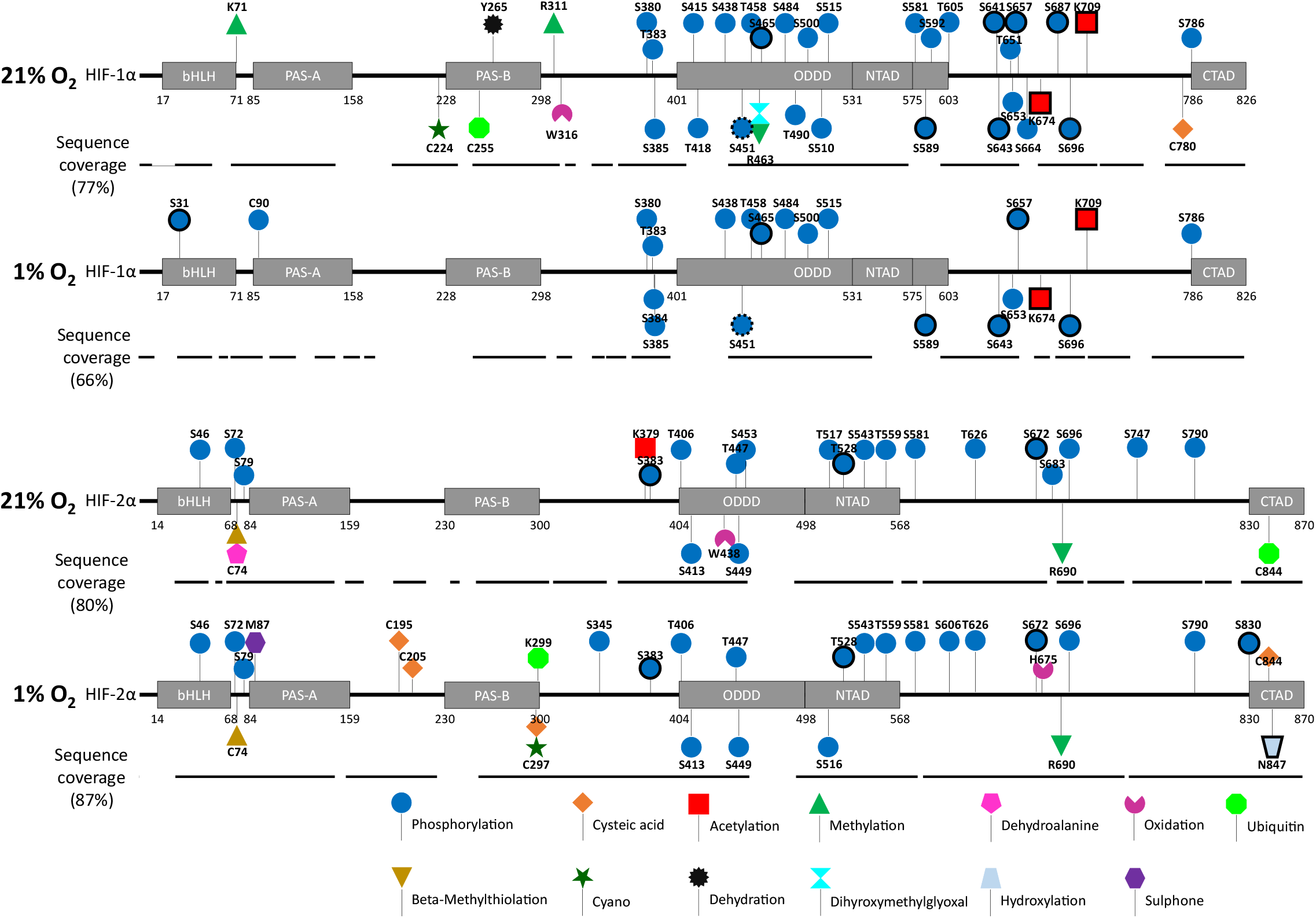
PTM sites identified on HA-Clover-HIF-1α and HA-Clover-HIF-2α expressed in HeLa cells in either 21% O_2_ or 1% O_2_. Schematic maps are presented detailing the major HIFα domains: bHLH=basic helix-loop-helix; PAS-A/B=Per/ARNT/Sim A/B; ODDD=O_2_ dependent degradation domain; NTAD=N-terminal transactivation domain; CTAD=C-terminal transactivation domain. All phosphorylation sites were identified using the MASCOT node within ProteomeDiscover and site localized using ptmRS (filtered to 1% FDR, MASCOT Score>20, PSMs>1 and ptmRS score>98.0). All other PTMs were identified using PEAKS PTM (filtered to 1% FDR, −log_10_*P*>30.0, Ascore>30, observed in both biological replicates for a given condition), n=2. Protein sequence coverage was determined from the culmination of different protease digests across all non-TiO_2_ enriched experiments, and is displayed below each schematic. Symbols with solid borders represent PTMs that have previously been identified in at least 1 LTP (low-throughput) study, symbols with dotted borders in at least 5 HTP (high-throughput) studies, according to PhosphositePlus (Hornbeck et al. 2012).

Our analyses revealed that both HIF-1α and HIF-2α are extensively decorated by PTMs with a total of 41 and 39 PTMs confidently mapped respectively to the two isoforms, encompassing 13 different types of modification (Fig. 5; Supp. Table 8). Comparison with publicly available PTMs information in UniProt (Consortium 2018) and PhosphositePlus (Hornbeck et al. 2012), suggests that >80% of our mapped PTMs are novel. Of the 37 and 15 sites identified in low-throughput (LTP) studies in PhosphositePlus (accessed 29^th^ July 2020) for HIF-1α and HIF-2α, respectively, we failed to obtain evidence for ~80% of them. This discrepancy is likely due to the methodology used to define these PTMs, having been largely elucidated using *in vitro* assays, compared to our *in cellulo* approach employing full length proteins. It is also worth noting that many of the previous cellular based studies exploited hypoxia mimicking drug treatments which are known to induce additional stress responses, as well as different cell lines, and thus are likely contribute to the observed PTM differences (Bracken *et al.* 2006, Tian *et al.* 2011, Ohyashiki *et al.* 2009 & Yu *et al.* 2011).

Over 50% of the PTMs that we mapped on the HA-Clover-HIFα isoforms, expressed in HeLa cells, were differentially observed at different O_2_ tensions. The PTM landscape of HIF-1α in particular was markedly different; of the 41 PTMs identified, 20 were specific to 21% O_2_ and three were observed only in 1% O_2_. However, it is worth noting that the ~10-fold overexpression of HA-Clover-HIF-1α in these experiments means that the biological relevance of the PTMs identified, particularly under normoxic conditions where HIF-1α would ordinarily be rapidly degraded, will need to be carefully evaluated. However, interestingly, the majority of the PTMs mapped as 21% O_2_ specific, localize to the O_2_-dependent degradation domain (ODDD), suggesting that these PTMs may have a role in the O_2_-dependent degradation of HIF-1α. Likewise, the PTM status of HIF-2α is also highly influenced by O_2_ tension. Of the 39 PTMs identified, 21 were O_2_-dependent with 8 being found only at 21% O_2_, and 13 being specific to 1% O_2_. However, unlike HIF-1α, these O_2_-dependent PTMs were distributed throughout the whole HIF-2α protein sequence rather than being more densely localized within the ODDD and the self-inhibitory domain (which lies between the N- and C-terminal transactivation domains; NTAD and CTAD) (Pugh *et al.* 1997 and Jiang *et al.* 1997).

Inspection of all PTMs across the two isoforms reveals that the main HIFα functional domains have highly different PTM maps. While the HIF-1α ODDD is hyper-phosphorylated (15 sites in total, 7 of which are O_2_-dependent), 6 PTMs were confidently localized to the HIF-2α ODDD, only two of which were O_2_-dependent. In contrast, while 5 different phosphorylation sites were localized to the N-terminal transactivation domain (NTAD) of HIF-2α (2 of which were O_2_-dependent), no PTM sites were identified in the HIF-1α NTAD. Although the inhibitory domain of both HIFα isoforms was densely populated with PTMs (accounting for ~25% of all PTMs for both isoforms), comparison of all identified sites of modification in this region, or indeed across the entirety of the two protein sequences, revealed limited PTM site correlation across the two isoforms. The relatively poor evolutionary conservation in the inhibitory domain between these two HIF isoforms, including a 60 residue insertion in HIF-2α (Supp. Fig. 3), raises the possibility that this region may act as an evolutionary ‘hot-pocket’ for evolving different/novel functions between isoforms, through PTM, which could explain some of the differences observed.

Among the range of PTMs detected, we specifically noted multiple types of cysteine modification. Due to the reactivity of the thiol side chain, cysteine can be subject to a variety of covalent modifications, notably by oxidative modification (making Cys a prime target for mediating redox-regulation of proteins), phosphorylation or ubiquitination (Chung *et al.* 2013, Sun *et al.* 2012, Hardman *et al.* 2019 and McDowell *et al.* 2013). Although some cysteine PTMs are readily reversible by reduction, *e.g.* during sample preparation with agents such as DTT (as used here), others are irreversible, or require catalytic activity of specific enzymes. Using PEAKS-PTM, we defined 6 different types of Cys modification (identified in both bioreps), including oxidation to cysteic acid (sulfonic acid), ubiquitination, cyanoylation, dehydroalanine, beta-methylthiolation and phosphorylation, across 12 different Cys residues, with 11 Cys-PTMs being O_2_-sensitive, 4 on HIF-1α, and 7 that were O_2_-sensitive on HIF-2α. Considering these findings in more detail, and the fact that the thioester linkage of Cys-Ubiqitin is likely susceptible to reduction (and thus loss) upon DTT treatment (Carvalho *et al.* 2007), we cannot rule out the possibility that the two sites of Cys-ubiquitination (Cys255 on HIF-1α and Cys844 on HIF-2α) may in fact be di-carbamidomethylation, which, with a delta mass of 114.0429 Da, is isobaric with the di-Gly remnant following tryptic proteolysis of a covalently linked ubiquitin (Kim *et al.* 2016). However, while beta-methylthiolation (R-S-S-CH3) should also be susceptible to DTT-mediated reduction, we were unable to identify alternatives to this modification when considering the delta mass of 45.9877 amu.

Of particular interest, we also identified a novel site of Cys phosphorylation, pCys90, on HIF-1α. Phosphorylation is typically associated with Ser, Thr and Tyr residues in mammalian proteins. However, multiple other residues in human proteins, including Cys, can also undergo phosphorylation. Indeed, Hardman *et al.* identified 55 sites of pCys in HeLa cell lysate proteins (at 1% FDR, ptmRS score >0.90). Manual spectral interpretation of the Cys90-containing phosphopeptide supported the likelihood of this phosphate group residing on the Cys residue (even though this peptide did not contain any putative canonical sites phosphorylation *i.e.* lacking Ser, Thr or Tyr) due to identification of y^4^, y^5^ y^6^, y^8^, y^9^ and y^10^ site-determining ions (Supp. Fig. 4), confirming pCys90 as a novel site of non-canonical phosphorylation on HIF-1α.

It is important to note that amongst the numerous modifications that we observed, there was no evidence of Pro hydroxylation. The absence of detection of hydroxyproline is likely due to the inherent rapid HIFα protein degradation that is triggered: t1/2 for HIF-1α is reported to be ~4-6 min in a manner dependent on Pro hydroxylation within the ODDD (Jaakkola *et al.* 2001, Ivan *et al.* 2001 and Moroz *et al.* 2009). We identified Asn hydroxylation, but only on HIF-2α and at 1% O_2_. These observations are likely explained by the fact that Asn hydroxylation has recently been shown to be reversible (Rodriguez *et al.* 2020), and that hydroxy-Asn containing HIFα molecules are likely additionally hydroxy-Pro and thus degraded, indirectly limiting their abundance.

### Hypoxia-induced phosphorylation of the highly conserved Ser31 on HIF-1α abrogates transcriptional activity without affecting protein stability

Conservation analysis (Supp. Table. 9) revealed that Ser31 within the bHLH domain of HIF-1α, which we identified as being phosphorylated in hypoxia only, is highly conserved amongst all species analyzed except Bony Fish (*Osteichthyes*), where the equivalent residue (in 72 out of the 76 *Osteichthyes* species) is a non-phosphorylatable glycine residue. Interestingly, it is only the more evolutionary distant (early) species of *Osteichthyes* that have a serine residue at this position, including Gar species, gray bichir and coelacanth, all of which pre-date the genome duplication of *Osteichthyes* (Betancur *et al.* 2013). We hypothesised therefore that Ser31 phosphorylation could play an important function in hypoxic-mediated transcriptional regulation of HIF-1α that is not required in most Bony Fish. Although Ser31 phosphorylation has previously been identified as a target for PKA using an *in vitro* assay on a recombinant fragment of HIF-1α, pSer31 has not previously been identified in cells on full length protein, and the role of this modification has not been defined beyond an absence of an effect on the stability of HIF-1α protein fragments (Bullen *et al.* 2016). Given the evolutionary distinct nature of Ser31, we set out to investigate the functional role of this modification in cells expressing full-length HIF-1α.

Modelling of Ser31 phosphorylation using the crystal structure of the HIF-1α/HIF-1β heterodimer bound to the HIF response element (HRE) (PDB: 4ZPR; Wu *et al.* 2015) with PyMol, predicted a distance of 9.3 Å between the phosphate group and the negatively charged DNA backbone (pyTMs plugin, Warnecke *et al.* 2014; Fig. 6A). Such close proximity of pSer31 to the DNA backbone suggested a possible role for this phosphorylation event in preventing DNA binding through charge repulsion, ultimately repressing transcription.

**Figure 6:**
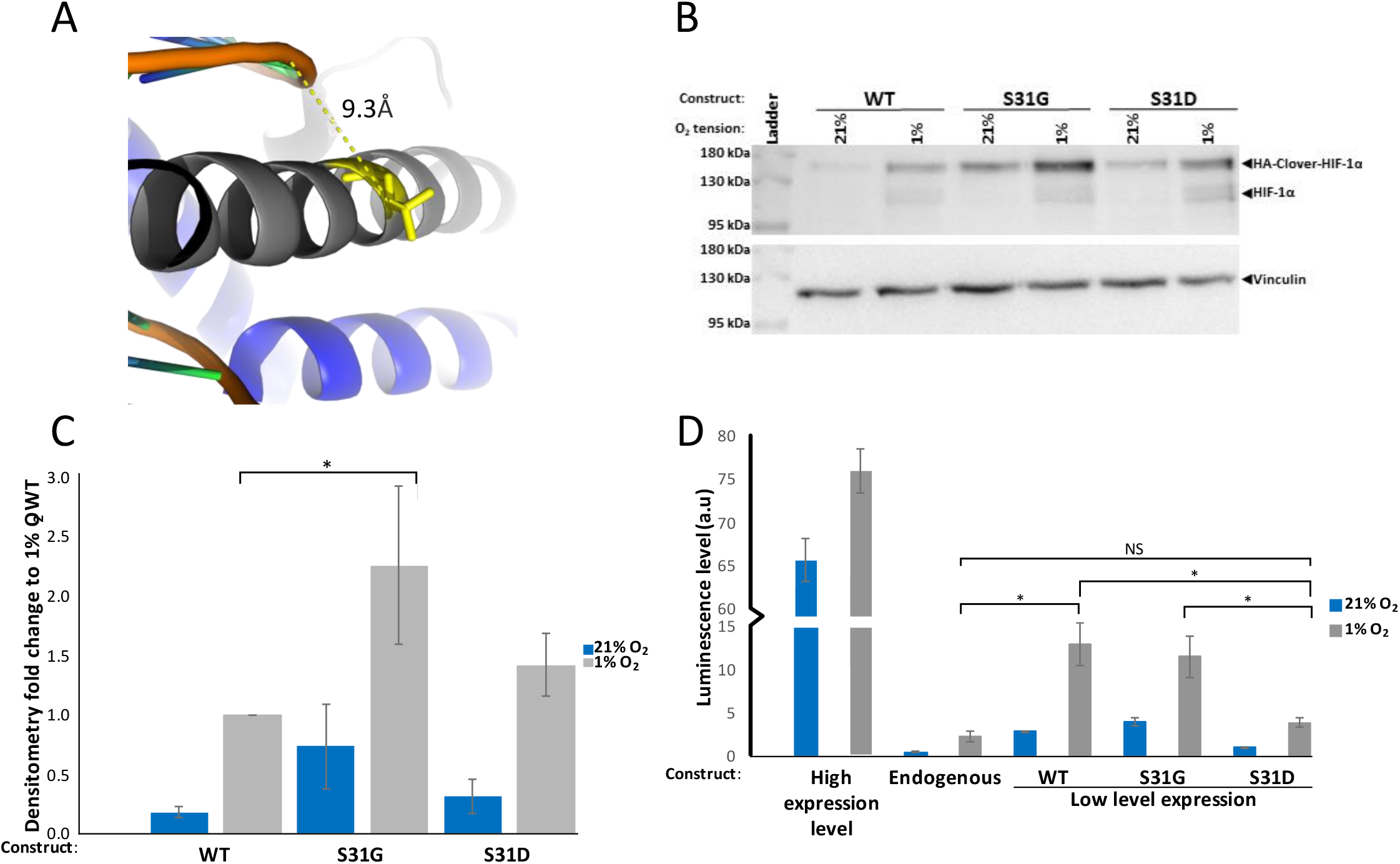
Biochemical characterization of HIF-1α Ser31 phosphorylation by mutational analysis. A) PyMol model of the HIF-1α/HIF-1β heterodimer bound to the hypoxia responsive element (HRE) (PDB:4ZPR) in silico phosphorylated at Ser31. Grey= HIF-1α, blue=HIF-1β, yellow=pSer31, orange=DNA backbone. Figure shows the shortest distance from the phosphate to DNA backbone measured as 9.3 Å. B) Western blot of HeLa cells expressing either wild-type (WT), phosphonull (S31G) or phosphomimetic (S31D) HIF-1α using an anti-HIF-1α (top) or anti-vinculin (bottom) antibody as loading control. Representative of n=3. C) Densitometry analysis of HIF-1α levels as determined in (B). Average fold-change (n=3) is shown +/− SEM) in response to 1% O_2_, normalized to vinculin protein levels. 2-sample t-tests were performed, *=*P*-value<0.05. Blue=21% O_2_, grey=1% O_2_. D) Luciferase reporter assay using HRE-Luciferase in presence of WT, S31G or S31D HIF-1α in HeLa cells at either a 1:1 ratio HA-Clover-HIF-1α:pcDNA(-) (high expression, as used for the IP/MS analysis, WT only), or low level expression (1:19 ratio) for all constructs. Average luminescence levels (from n=3) is reported +/− SEM. Blue=21% O_2_, grey=1% O_2_. 2-sample t-tests were performed, *=*P*-value<0.05, NS=not significant.

To investigate the impact of HIF-1α Ser31 phosphorylation on transcriptional activity and stability, we expressed either wild-type (WT) HA-Clover-HIF-1α, a phosphonull form of HIF-1α where Ser31 was replaced with Gly (HA-Clover-HIF-1α S31G), as present in bony fish, and a phosphomimetic version replacing Ser31 with Asp (HA-Clover-HIF-1α S31D). Over-expression of S31G HIF-1α, resulted in a significant two-fold increase in protein stability in 1% O_2_ (p <0.05, n=3; Fig. 6B/C), while there was no statistically significant difference in protein stability for S31D HIF-1α, or for either construct under 21% O_2_.

Using an HRE-luciferase reporter as a proxy to investigate HIF-dependent transcription, we next evaluated hypoxia-dependent regulation of transcription in cells from endogenous HIF proteins, and following transfection of WT, S31G or S31D HIF-1α. Despite the ~2-fold increase in expression level of S31G HIF-1α compared to WT (Fig. 6B/C), there was no noticeable difference in transcriptional activity (Fig. 6D) under either normoxic or hypoxic conditions. However, in agreement with our hypothesis, we did observe a significant (*p* < 0.05) ~5-fold reduction in HRE transcriptional output with the S31D phosphomimetic compared to WT-, or S31G-HIF-1α. Indeed, the transcriptional output from S31D approaches that observed with endogenous HIF proteins in both 21% and 1% O_2_, suggesting that this mutant is transcriptionally inactive, and HIF-1α Ser31 phosphorylation functions as a transcriptional repressor. Thus, this additional regulation of HIF is lost in the bony fish.

### Phosphorylation of HIF-2α adjacent to LAP proline hydroxylation motifs affects both protein stability and transcriptional output

The canonical Pro hydroxylation sites on HIF-1α (Pro402 and Pro564) and HIF-2α (Pro405 and Pro531), which lie in the consensus motif LXXLAP (Ivan *et al.* 2001, Jaakkola *et al.* 2001 and Masson *et al.* 2001), localize to regions that are highly conserved both between the two HIF isoforms (Fig. 6A), and throughout evolution (Loenarz *et al.* 2011) emphasizing their essential roles in regulating HIF protein function. However, there is some flexibility in this Pro hydroxylation motif, notably at the −5, −2, and −1 positions relative to the hydroxy-Pro residue (Huang *et al.* 2002). Given this flexibility, we hypothesize that the mammalian specific (FXXLAP576) motif on HIF-2α could constitute a third, yet unidentified as being hydroxylated, LAP motif (Fig. 6A) that may have functional roles.

The observation that both known HIF-2α Pro hydroxylation sites are adjacent to Thr residues, which we have identified as being phosphorylated (pThr406 and pThr528), with an additional site of phosphorylation identified at Ser581, immediately downstream of the putative Pro576 hydroxylation site, raises the possibility that there may be interplay between phosphorylation of these residues and hydroxy-Pro, and thus (indirect) regulation of HIF-2α stability. All three phosphorylation sites on HIF-2α are oxygen-independent (Fig. 4) and are highly conserved (Supp. table 9). In contrast, HIF-1α does not have these Thr residues at the equivalent positions within its canonical hydroxy-Pro sites, rather having non-phosphorylatable Ala or Met residues (HIF-1α: Ala403 and Met561). Additionally, the putative HIF-2α Pro576 site is in a highly divergent region between isoforms, indicating that the roles of these pThr/pSer are HIF-2α specific. Using phosphonull (T406A, T528A, S581A) and phosphomimetic (T406E, T528E, S581D) versions of HIF-2α, we evaluated protein stability and transcriptional activity in HeLa cells as above (Fig. 7).

**Figure 7:**
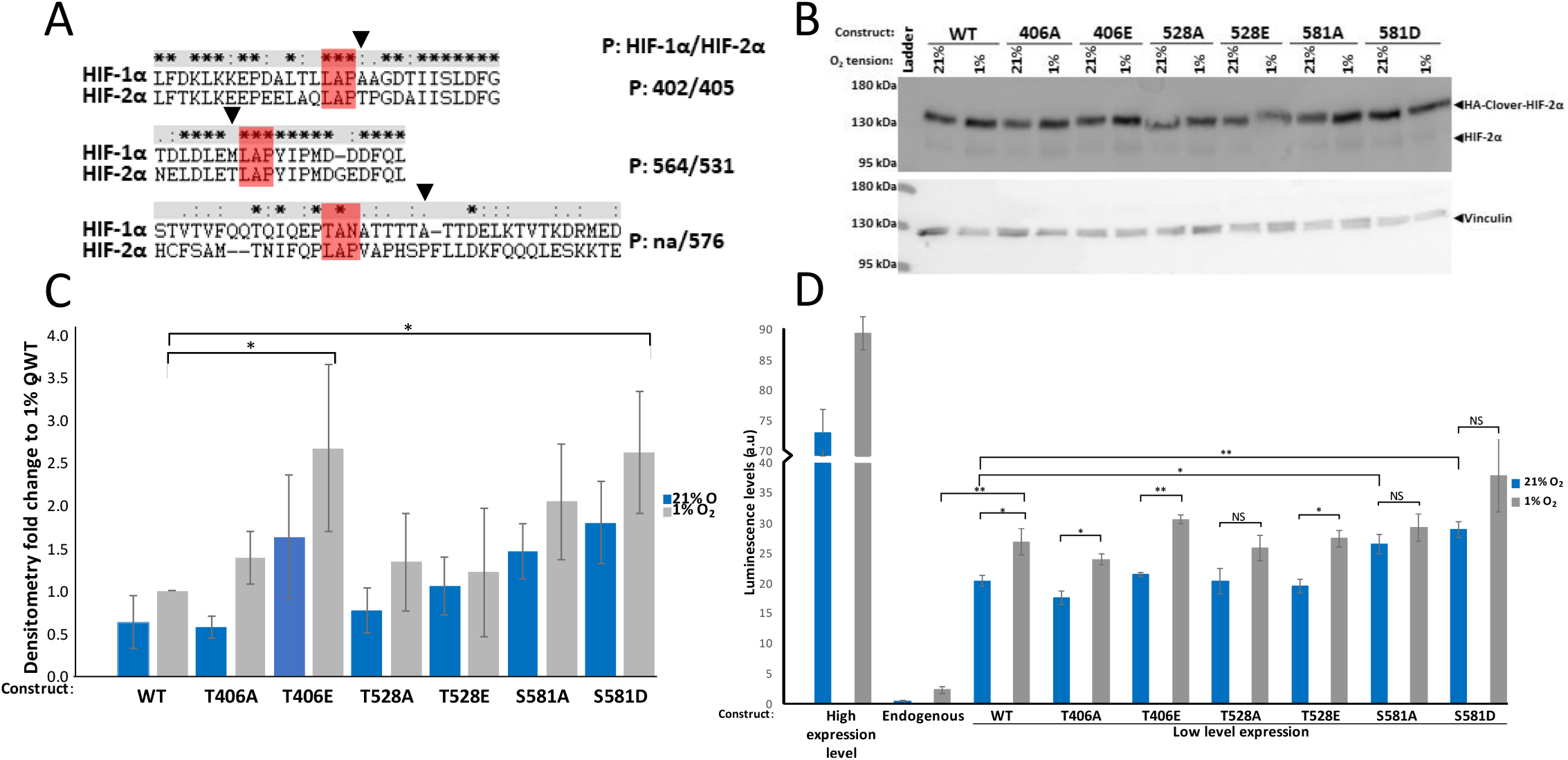
Biochemical characterization of HA-Clover-HIF2α Thr 406, Thr528 and Ser581 phosphorylation by mutational analysis. A) Sequence alignments of HIF-1α and HIF-2α surrounding the canonical proline hydroxylation sites Pro402/Pro564 and Pro405/Pro531 respectively) and a 3^rd^ putative LAP motif (none, Pro576 respectively). Alignments were performed using the MUSCLE multiple sequence alignment tool and viewed in Clustal X, * = identical residue: = similar residue properties. = less similar residue properties space = no similarity in residues (properties as determined by Clustal X) -= gap inserted to align sequences. Triangles identify HIF-2α specific phosphorylation sites, red highlights the putative LAP motif. B) Western blot of HeLa cells expressing either wild-type (WT), phospho-null (Thr to Ala) or phospho-mimetic (Thr to Glu, or Ser to Asp) HIF-2α using an anti-HIF-2α (top) or anti-vinculin (bottom) antibody as loading control. Representative of n=3. C) Densitometry analysis of HIF-2α levels as determined in (B). Average fold-change (n=3) is shown +/− SEM) in response to 1% O_2_, normalized to vinculin protein levels. 2-sample t-tests were performed, *=*P*-value<0.05. Blue=21% O_2_, grey=1% O_2_. D) Luciferase reporter assay using HRE-Luciferase in presence of WT, phospho-null or phospho-mimetic HIF-2α in HeLa cells at either a 1:1 ratio HA-Clover-HIF-2α:pcDNA(-) as used for the IP/MS analysis (high expression levels, WT only), or low level expression (1:19 ratio) for all constructs. Average luminescence levels (from n=3) are reported +/− SEM. Blue=21% O_2_, grey=1% O_2_. 2-sample t-tests were performed, *=P-value<0.05, NS=not significant.

Compared to HA-Clover-HIF-2α WT controls, protein levels of T406E HIF-2α were elevated ~2.5 fold at 1% O_2_ (*p*-value <0.05; Fig. 6B/C), although this was not associated with an increased transcriptional activity (Fig 6D). There was no significant effect on either stability or transcriptional function for T406A (at either 21% or 1% O_2_), suggesting that Thr406 phosphorylation acts to stabilize HIF-2α protein levels in an O_2_-independent manner. In agreement with Pangou *et al.* 2016, we did not observe any significant effect on protein stability for either of the Thr528 mutations (Fig. 6B/C). However, while Pangou *et al.* 2016 report a ~50% decrease in the transcriptional activity of T528A HIF-2α, due to reduced nuclear localization, we did not observe any changes in transcriptional output for either of the Thr528 mutations (Fig. 6D). Mutation of Ser581 to either Ala or Asp (S581A or S581D respectively) increased protein stability in an oxygen-independent manner by > 2-fold, although was only statistically different for S581D in 1% O_2_. Similarly to Thr406 and Thr528 mutations, neither mutation of Ser581 lead to transcriptional effects in 1% O_2_. However, interestingly, the transcriptional output of both Ser581 mutants in 21% O_2_ were elevated, reaching a statistically significant difference compared to WT, and were no longer statistically different to the respective 1% O_2_ readings, *p*<0.05). Hence, this suggests a potential mechanism to activate HIF-2α signaling by phosphorylation in oxygenated conditions. As both phospho-mimetic and phospho-null mutants of Ser581 result in elevated transcriptional activity at 21% O_2_, we suggest a potential dual role of Ser581, maybe in tandem with the putative hydroxy-Pro576 site, where phosphorylation promotes HIF-2α dependent transcription by prevention of a 21% O_2_ specific inhibitory pathway.

## DISCUSSION

Here we present the first unbiased discovery study to characterize the regulation of full-length cellular HIF-1α and HIF-2α by PTM, and to explore changes in their protein interaction networks as a function of oxygen tension. The extremely low levels of these proteins and their rapid degradation under normoxic conditions, necessitated overexpression (~10-fold compared to endogenous protein levels) of tagged (HA-Clover) HIF-1α or HIF-2α (Supp. Fig. 1A/B) for sensitive PTM mapping. However, even under these conditions we demonstrated correct sub-cellular localization and hypoxia-induced increase in transcriptional activity for both proteins.

### Use of overexpression of HA-Clover-HIFα

The comparability of HA-Clover-HIF-1α protein levels at different oxygen concentrations, *i.e.* the significant reduction in proteolysis in normoxia, would suggest that an aspect of the degradation pathway was saturated. However, the overexpression of HA-Clover-HIF-1α at these levels did not affect the oxygen dependency of the endogenous HIF-1α protein (Supp. Fig. 1B), and given our lack of observation of sites of Pro-hydroxylation whilst identifying all three EGLN PHDs and the majority of the 26S proteasomal subunits as binding partners, we suggest that there may be an oxygen-dependent pathway which acts prior to the canonical proline hydroxylation pathway, yet fundamentally fails to act on the HA-Clover-HIFα constructs. One explanation could be the fact that endogenous HIFα mRNA contains a 5′ UTR TOP motif known to regulate HIFα translation in an oxygen-dependent manner, through the PI-3-K/mTOR/S6K pathway (Zhong *et al.* 2000 & Thomas *et al.* 2006), which is lost in the HA-Clover-HIFα cDNA-containing constructs.

### HIFα interaction with mitochondrial proteins

Our data show that the protein interaction networks of HIF-1/2α are highly influenced by oxygen tension with ~45-50% of (direct or indirect) protein binding partners being identified only under specific oxygen conditions. O_2_-dependent interactome regulation is particularly apparent for HIF-2α where ~7.5 times as many proteins were identified as hypoxia-specific binding partners (84 *vs* 642 proteins identified as 21% and 1% O_2_-specific respectively), with the levels of an additional ~380 proteins being significantly higher in hypoxia. In contrast, fewer unique or differentially regulated proteins were identified with HIF-1α under hypoxic conditions compared with 21% O_2_. Amongst the binding partners of both HIFα isoforms were a number of proteins associated with the mitochondria under all O_2_ conditions. While the association of HIFα and mitochondria cross-regulation is well known (Thomas *et al.* 2019), our findings support studies localizing a sub-population of HIFα proteins to this organelle (Briston *et al.* 2011 & Concolino *et al.* 2018), and suggest that a proportion of HIF-1/2α may be associated with mitochondria in a manner independent of oxygen tension. However, for HIF-2α, whilst most mitochondrial-associated interactors were O_2_-independent, by considering only O_2_-tension specific interactors, significantly more mitochondrial-associated proteins were identified as hypoxia specific (~7 fold), suggesting an oxygen-sensing role in HIF-2α sub-cellular localization, and potential additional roles for this isoform as a transcription factor of mitochondrial DNA.

Comparing the interactomes of HIF-1α and HIF-2α under normoxic conditions revealed minimal overlap; only 22 proteins were common between these two isoforms in the cohort of proteins that were specific to 21% O_2_. Combined with our quantification of ~4 fold more normoxic specific protein interactors for HIF-1α, our data suggest distinct regulatory mechanisms for HIF-1α and HIF-2α under physiological oxygen conditions. However, for both isoforms under hypoxic conditions, we observed upregulation of proteins associated with telomere regulation as well as components of the RAN signaling pathway, which is involved in nuclear export of the telomerase protein TERT. This is in line with a recent study in neuroblastoma, showing that the hypoxic tumours are associated with mechanisms of telomerase activation (Cangelosi *et al.* 2020). These findings, combined with the observation of numerous mitochondrial proteins (and mitochondrial dysfunction as an enriched pathway) is interesting in the context of the known relationship between mitochondrial dysfunction and telomere damage (Zheng *et al.* 2019), suggesting mechanistic roles for HIF-1/2α in hypoxia-dependent regulation of these processes.

### HIFα cysteine modifications

Our PTM data shows that both HIF-1α and HIF-2α are subject to extensive modification, with ~40 different sites of PTM (13 different types of PTM) confidently localized on both isoforms. Although these PTMs were distributed throughout the whole length of the two proteins, the majority of the sites that we identified cluster in the *C*-terminal region, particularly the ODDD and the self-inhibitory domains. Due to the analytical strategy employed (including phosphopeptide enrichment), it is perhaps unsurprising that the majority of PTMs identified in this study were due to phosphorylation. Over 80% of the PTMs that we identified are novel, including a site of Cys phosphorylation on HIF-1α (pCys90), which we observed under all conditions. Given the important role of Cys PTMs in regulating redox-sensitive protein functions, and the other Cys modifications that we identified here, it is likely that more (reversible) Cys modifications will be observed in conditions that do not employ a reducing agent during sample preparation.

### Transcriptional regulation of HIF by phosphorylation

To aid functional characterization of individual PTM sites, we used a combination of evolutionary, domain and isoform comparison analyses. We have identified a role for the highly evolutionarily conserved, hypoxia-induced, pSer31 in the bHLH domain of HIF-1α in abrogating transcription, as well as a putative role for pSer581 on HIF-2α in increasing the, usually low, transcriptional activity in normoxia. Based on observation of elevated levels of phospho-mimetic mutants compared with phospho-null or WT controls, HIF-2α phosphorylation on Thr406 and Ser581, but not Thr528, appear to prevent protein degradation in a manner that is largely independent of O_2_, although is only significant under hypoxia. While we hypothesize that these sites of phosphorylation, lying in close proximity to hydroxy-Pro residues (within the PHD LXXLAP consensus), may work in concert with Pro hydroxylation to regulate protein degradation, we were unable to observe Pro hydroxylation. In support of this theory, it is interesting to note that both IKKα and IKKβ contain LXXLAP motifs (with Pro191 of IKKβ being a known hydroxylation site, Cummins *et al.* 2006) that lie in close proximity to regulatory phosphorylation site. However, there is currently no clear evidence for cross-talk between these PTMs and will require future investigations in presence of proteasome inhibition.

## CONCLUSIONS

We have demonstrated that the oxygen-dependent signaling of HIF, through extensive PTM of the HIF-1/2α subunits and significant changes in the HIF interactomes, is significantly more complicated than was previously appreciated. These data will thus help to delineate the different physiological roles of these closely related isoforms and can be used to start to unravel their mechanistic involvement in fine-tuning the hypoxic response.

## MATERIALS and METHODS

### Reagents

Tissue culture reagents were purchased from Gibco (ThermoFisher Scientific). Powdered chemical reagents and custom DNA primers were purchased from Sigma-Aldrich. Mass spectrometry (MS) solvents were purchased from ThermoFisher Scientific, HPLC grade. All Eppendorf tubes used for MS are Ultra-High recovery Eppendorf tubes (STARLAB).

### Cloning of HA-Clover plasmids

All cloning was performed using the In-Fusion Cloning technology (TakaraBio). Restriction enzymes were purchased in their HF format from New England Biolabs (NEB); PCR stages used the KOD HotStart Polymerase Kit (Sigma-Aldrich); In-gel DNA extraction steps used the E.Z.N.A gel extraction kit (omega BIO-TEK). Plasmid amplification was done using the PureLink HiPure Plasmid Maxiprep Kit (Invitrogen). All cloned plasmids were validated by LightRun sequencing (Eurofins genomics). A HA-HIF-1α plasmid was obtained as a gift from Prof. Sonia Rocha, University of Liverpool, UK. To generate HA-Clover-HIF-1α, HA-HIF-1α was linearized with BamHI and had the clover gene amplified from the pcDNA3-Clover plasmid (Addgene #40259). To generate HA-Clover, HA-HIF-1α had the HIF-1α gene removed from HA-HIF-1α by double restriction digestion with BamHI and EcoRV. The clover gene was amplified as before. To generate HA-Clover-HIF-2α, the HA-Clover plasmid was linearized with BsrGI, and the HIF-2α gene amplified from a HaloTag-HIF-2α plasmid (Kazusa DNA Research Institute #pFN21AB4384). The newly generated HA-Clover-HIF-1α and HA-Clover-HIF-2α are available via Addgene.

### Site Directed Mutagenesis

The ‘MEGAprimer’ technique was used to create all mutants (Sarkar *et al.* 1990). Briefly, a 1^st^ round PCR using a flanking primer and a mutagenic primer was used to create a PCR fragment with the desired mutation. In a 2^nd^ round, the PCR mutagenic fragment was used as a primer with a second flanking primer to synthesize the full-length gene, which is re-inserted into the plasmid backbone through restriction digestion and ligation. Flanking primers (used for mutagenesis of either HA-Clover-HIF-1α / 2α): Forward 5′ GATCCGCCACAACGTTGAG 3′ and Reverse 5′ AGACAATGCGATGCAATTTCC 3′. Mutagenic primers were designed to create the smallest PCR fragment in the 1^st^ round PCR. Mutagenic primers: S31D 5′ CAGATTCTTTACCTCGCCGAG 3′, S31A 5′ CAGATTCTTTATCTCGCCGAG 3′, T406E 5′ GTCTCCTGGTTCGGGAGCCAG 3′, T406A 5′ CTCCTGGGGCGGGAGCC 3′, T528E 5′ CTTGGAGGAACTGGCACC 3′, T528A 5′ CTTGGAGGCACTGGCACC 3′, S581D 5′CCCGCACGATCCCTTCC 3′, S581A 5′ CCCGCACGCTCCCTTCC 3′. Restriction digestion was performed with BsrGI and NotI for all plasmids and full-length PCR products.

### Cell Culture, Transient Transfection and Hypoxic Treatment

HeLa cells (ECACC catalogue #: 93021013) were seeded at a density of ~1.75 x 10^5^ cells/cm^2^ in DMEM media supplemented with 10% (v/v) foetal calf serum, 1% (v/v) non-essential amino acids and 1% (v/v) penicillin/streptomycin and incubated at 37 °C, 5% CO_2_, 21% O_2_ for 24 h. Cells tested for mycoplasma infection monthly. Transient transfection was performed 24 h prior to experimental use, using PEI 40K MAX, Linear (PEI, Polysciences). Plasmid stocks were diluted with unsupplemented DMEM to a final DNA concentration of 10 ng/μl. For immunoprecipitation and mass spectrometry experiments (high expression levels), a ratio of 1:1 HA-Clover plasmid:pcDNA(-) (Invitrogen #V79520) DNA was used. For western blotting experiments a ratio of 1:19 was used, and for luciferase assays a ratio of 1:9:10 HA-Clover plasmid:pcDNA(-):HRE-Luciferase (Addgene #26731) was used (physiological expression levels). A stock 1% PEI (w/v) in PBS (pH 7.5, adjusted with NaOH) solution was used at a ratio of 4 μL:1 μg total DNA, vortexed briefly and incubated at room temperature for 30 min. The equivalent of 5% of the total cell culture volume of transfection mixture was added to cells, and incubated for 18 h. The cell media was then replaced by fresh growth media and incubated in either 21% O_2_ or 1% O_2_ in a H35 hypoxic station (Don Whitley Scientific) for 4 h prior to lysis.

### Immunoprecipitation and Sample Preparation for Proteomics

PBS and lysis buffer (50 mM Tris-HCl pH 8.0, 120 mM NaCl, 5 mM EDTA, 0.5% (v/v) Nonidet P-40, 1X EDTA-free cOmplete protease inhibitor (Roche) and 1X phosSTOP (Roche)) were pre-conditioned identically to experimental conditions for cellular incubation, 4 h at either 21% or 1% O_2_, prior to use. Cells (~1.5 x 10^8^) were washed in an equal volume of PBS and lysed in 12 μL of lysis buffer per 100,000 cells, scraped and combined into a 50 ml Falcon tube (Eppendorf) before removal of the hypoxic samples from the hypoxystation. Lysates were rotated for 30 min at 4 °C before centrifugation at 10,000 *g* for 10 min at 4 °C, and collection of the clarified supernatant. To reduce the final concentration of Nonidet P-40 to 0.2% (v/v), the supernatant was then diluted 2.5X in dilution buffer (lysis buffer lacking detergent). Unless otherwise stated, all steps were performed at 4 °C and centrifugation steps were performed at 3,000 *g* for 2 min at 4 °C. All beads were washed 3 times in 5 volumes of dilution buffer before use. Diluted lysate was pre-cleared by incubation with binding control agarose beads (bab-20, CHROMOTEK), at a volume of slurry equivalent to 1:200 (v/v) of bab-20 beads:diluted lysate, rotating end-over-end (1 h, RT) before centrifugation and collection of supernatant. Immunoprecipitation was performed by incubation with GFP-TRAP_A beads (CHROMOTEK) at a volume of slurry equivalent to 1:800 (v/v) GFP-TRAP_A beads:diluted lysate and rotating end-over-end for 18 h at 4 °C. Bound complexes were washed 3 times in 5 volumes of dilution buffer, then twice in 25 mM HPLC grade ammonium bicarbonate (AmBic). Proteins were eluted in 1% (v/v) RapiGest SF (WATERS) in AmBic, volume equal to that of GFP-TRAPS_A bead slurry used. Eluant was heated at 95 °C for 15 min, vortexing for 5 s every 2.5 min. Supernatant was collected following centrifugation (10,000 *g*, 5 min, RT) and diluted with AmBic to achieve a final RapiGest concentration of 0.06%. Protein concentration was determined using a Nanodrop 2000 at a wavelength of 205 nm. Proteins were then subject to reduction with DTT and alkylation with iodoacetamide as previously described (Ferries *et al.* 2017). The eluent was equally divided into three for digestion with either: 10:1 (w/w) Trypsin Gold (Promega), 7.5:1 (w/w) Chymotrypsin (Promega) or 5:1 (w/w) Elastase (Promega), using manufacturer recommended temperatures for 18 h with 600 rpm shaking. Rapigest SF was hydrolysed and removed from samples as described by Ferries *et al.* 2017. Digests were subjected to in-house packed, strong cation exchange stage tips (Empore™ Supelco 47 mm Cation Exchange disc #2251, 3 discs/200 μL tip). All centrifugation steps were at 2000 g for 4 min (or until all liquid had passed through the tip) at RT, tips were equilibrated by two sequential washes with 200 μL of each: acetone, methanol, water, 5% (v/v) ammonium hydroxide (in water) and water, before addition of a peptide sample to the stage tip. A wash was performed with 250 μL of 1.5% (v/v) TFA and elution was done with 250 μL of 5% ammonium hydroxide (in water). Samples were split 95%:5% and dried to completion under cooled vacuum centrifugation. 95% samples were subjected to Titanium dioxide (TiO_2_) phosphopeptide enrichment, as described by Ferries *et al.* 2017, solubilizing peptides in a volume of 100 μl. All dried peptides were solubilized in 20 μL of 3% (v/v) ACN, 0.1% (v/v) TFA in water, sonicated for 10 min and centrifuged 13,000 g for 15 min at 4 °C prior to liquid chromatography-mass spectrometry analysis (5%: High-Low method, 95%: High-High method).

### Liquid Chromatography and Mass Spectrometry Analysis

Peptides were separated using an Ultimate 3000 nano system (Dionex) by reversed phase HPLC, over a 60 min gradient, as described by Ferries *et al.* 2017. All data acquisition was performed using a Thermo Orbitrap Fusion Tribrid mass spectrometer (Thermo Scientific), with HCD fragmentation set at 32% normalized collision energy (NCE) for 2+ to 5+ charge states. For 5% samples a High-Low method was used, MS1 spectra were acquired in the Orbitrap (60K resolution at 200 *m/z*) over a range of 350-2000 *m/z*, AGC target = 2e^5^, maximum injection time = 100 ms, with an intensity threshold for fragmentation of 5e^4^. MS2 spectra were acquired in the Iontrap set to rapid mode (15K resolution at 200 *m/z*), maximum injection time = 50 ms with a 1 min dynamic exclusion window applied at a 0.5 Da mass tolerance. For phosphopeptide enriched samples a High-High method was used, MS1 spectra were acquired in the Orbitrap (60K resolution at 200 *m/z*) over a range of 350-2000 *m/z*, AGC target = 5e^5^, maximum injection time = 250 ms, with an intensity threshold for fragmentation of 2e^4^. MS2 spectra were acquired in the Orbitrap (30K resolution at 200 *m/z*), maximum injection time = 250 ms and 2 μ-scans were performed. A dynamic exclusion window of 5 s was applied at a 10 ppm mass tolerance.

### Mass Spectrometry Data Analysis

Digested samples were analyzed through PEAKS Studio (v10.0) and MaxQuant (viewed in Perseus) v1.6.7.0, in conjunction with Andromeda. All database searches were against the UniProt Human reviewed database (updated April 2020), and the inbuilt contaminant database for MaxQuant. Enzyme parameters for all software (cleavage pattern: maximum number of permitted miscleaves): Trypsin (K/R –not P: 2), Chymotrypsin (F/Y/W/L: 4), Elastase (A/V/S/G/L/I: 8). PEAKS Studio settings: Instrument = Orbi-Trap, Fragmentation = HCD, acquisition = DDA, De Novo details = standard, variable modifications = carabamidomethylation (C) and oxidation (M) with a maximum number of 5 variable PTMs per peptide, MS1 mass tolerance = 10 ppm, MS2 mass tolerance = 0.5 Da. PEAKS PTM mode was enabled with a De novo score threshold of 15, identifications were filtered to a −log_10_P-value and Ascore >30.0. MaxQuant settings: split peaks was disabled and peptides for identification were only selected if 7 amino acids or longer, constant carbamidomethylation (C) and variable modifications of oxidation (M) and N-term acetylation. Default instrument parameter settings were used for an Orbitrap-Iontrap system, match between runs was enabled with a time window of 10 min. MS1 and MS2 tolerances were as used for PEAKS Studio. Perseus was used to group peptides by unique protein identifier and estimate protein abundance by averaging peptide intensities. Data was both filtered for identifications that were observed in a minimum of both replicates of an oxygen condition, or had missing values imputed with a normalized distribution below the lowest intensity peptides identified for label free quantification (LFQ). LFQ data had each replicate normalized to the level of HIF-1α/2α for that replicate. A 2-sample t-test was performed between oxygen conditions, s0 = 0.1, *p*-value correction = FDR based permutations (250 replicates). Data was exported and 1% protein intensities / 21% protein intensities calculated, any 0 value *q-*values were generated by dividing the lowest *q-*value by 10. Fold change (1%/21%) and q-values were transformed (log_2_) and imported into a custom R script.

Phosphopeptide enriched samples were analyzed using PEAKs studio as above (using the UniProt Human reviewed database (updated April 2020)), except for the following adjustments: instrument = Orbi-Orbi, additional variable modifications of phospho S/T/Y and MS2 mass tolerance = 0.01 Da. Proteome Discoverer (PD) in conjunction with MASCOT v2.6 was also used to analyze phosphopeptide enriched data (according to Ferries *et al.* 2017 & Hardman *et al.* 2019). A custom database was used for PD analysis of phosphopeptides, as described by Hardman *et al.* 2019, which was generated from all identifications from the respective trypsin digested (5%) sample that was analyzed by PD on the UniProt Human reviewed database (updated April 2020) with fixed modification = carbamidomethylation (C), variable modification = oxidation (M), instrument type = ESI-FTICR, MS1 mass tolerance = 10 ppm and MS2 mass tolerance = 0.5 Da. Phosphopeptide analysis PD settings: instrument type = ESI-FTICR, variable modifications = carbamidomethylaion (C), oxidation (M) and phospho (S/T/Y/D/E/K/R/H/C). MS1 mass tolerance = 10 ppm, MS2 mass tolerance = 0.01 Da and the ptmRS node on, filtered to a ptmRS score >98.0. All data from each analysis pipeline was filtered to a 1% false discovery rate (FDR) at the PSM level. Any phospho D/E/K/R/H/C sites identified were validated by manual spectra annotation.

### Phylogenetic analysis

Human HIF-1α (Q16665) and HIF-2α (Q99814) protein sequences were used as starting points to create phylogenetic trees following guidance from Hall 2013. Briefly, the selected protein sequences were BLAST searched for top 500 homologous sequences, a manual filter of 50% sequence homology was applied due to identification of the other HIFα isoform. All ‘partial’ and ‘unknown protein’ labelled proteins were removed. When multiple isoforms exist for a single species, reciprocal blast searching was performed and only the most similar to the human sequence was maintained in the dataset. Genus-species names were converted to ‘common names’ using the Taxize plugin for R (Chamberlain *et al.* 2013, using the Global Names Resolver (GNR) webpage [https://resolver.globalnames.org/]). Sequences were aligned using the multiple sequence alignment tool MUSCLE (Edgar 2004) and a phylogenetic tree was produced using MEGA7 (Kumar *et al.* 2016) with 500 bootstrap replicates.

### Functional enrichment analysis

DAVID Bioinformatics Resources (v6.8, Huang da *et al.* 2009 & Huang da *et al.* 2009) was used to understand isoform specific differences in HIF bound proteins as a function of O_2_ tension (*q*-value<0.05). Data was manipulated to maintain GOterm cellular compartment and GOterm biological process only and used in a custom R-script, *p*-value correction = Benjamini-Hochberg. LFQ data were analyzed through IPA software (QIAGEN Inc., https://www.qiagenbioinformatics.com/products/ingenuitypathway-analysis) to identify cellular pathways, and determine whether they are activated or inactivated by O_2_ tension.

### Western blot

Primary antibody incubations were performed for 18 h at 4 °C with the following antibodies: HIF-1α (Proteintech #20960-1-AP, 1/1000), and HIF-2α (Bethyl laboratories #A700-003, 1/1000), β-actin (Abcam #ab8226, 1/5000) and vinculin (Abcam #ab129002, 1/5000). Secondary antibody incubation was for 1 h at RT with: anti-rabbit HRP (cell signaling #7074s, 1/3000) or anti-mouse HRP (cell signaling #7076, 1/3000). Blots were visualized using ECL clarity (BIORAD) and images captured on a Syngene gel imaging G-Box.

### Luciferase assays

Post incubation in 21% or 1% O_2_, cells were washed in PBS and lysed in 25 mM Tris-phosphate pH 7.5, 15% (v/v) glycerol, 1% (w/v) BSA, 8 mM MgCl2, 0.1 mM EDTA, 2 mM DTT, 1% (v/v) Triton X-100 (60 μL/100,000 cells) and shaken for 5 min at room temperature. Lysate was split equally into 3 wells (80 μl per well) of a white 96 well plate (Greiner), and 200 μl of luciferin working solution added (500 μM Luciferin (Abcam #ab145164), 5 μM ATP (Sigma-Aldrich #FLAAS-1VL) in lysis buffer). Plates were incubated for 5 min in the dark, and endpoint luminescence measurements taken using a BMG Labtech FLUOstar Omega plate reader.

## SUPPLEMENTARY MATERIALS LIST

Supp. Fig. 1: Immunoprecipitation of transfected HA-Clover-HIFα.

Supp. Fig. 2: Hierarchal clustering of enriched pathways from HIF-1α and HIF-2α binding partners in different O_2_ conditions.

Supp. Fig. 3: Sequence alignment of the inhibitory domains of HIF-1α versus HIF-2^α^.

Supp. Fig. 4: HIF-1α pCys90 annotated MS2 spectrum.

Supp. File. 1: HIF-1α sequence alignment data.

Supp. File. 2: HIF-2α sequence alignment data.

Supp. Table. 1: Oxygen dependent protein binding partners of HIF-1α and HIF-2α.

Supp. Table. 2: Label free quantification of protein binding partners of HIF-1α and HIF-2α.

Supp. Table. 3: Gene ontology analysis of oxygen dependent HIF-1α and HIF-2α binding partners.

Supp. Table. 4: IPA pathway analysis of HIF-1α and HIF-2α binding partners

Supp. Table. 5: Oxygen-dependent protein binding partners of HIF-1α versus HIF-2α.

Supp. Table. 6: Gene ontology analysis of oxygen (in)dependent binding partners identified for both HIF-1α and HIF-2α.

Supp. Table. 7: IPA pathway analysis of oxygen (in)dependent binding partners identified for both HIF-1α and HIF-2α.

Supp. Table. 8: Mass spectrometry supporting data for HIF-1/2a PTM identification.

Supp. Table. 9: Previous identification and evolutionary conservation data of identified PTMs.

## Acknowledgements

We thank the technical support received for confocal imaging by the Centre for Cell Imaging staff, especially Jennifer Adcott for her support and assistance in this work. We thank Prof. Sonia Rocha (University of Liverpool, UK) for the gift of a HA-HIF-1α plasmid.

## Author Contributions

The manuscript was written through contributions of all authors. All authors have given approval to the final version of the manuscript.

## Data and materials availability

The mass spectrometry proteomics data have been deposited to the ProteomeXchange Consortium via the PRIDE partner repository with the dataset identifier PXD022479. Generated plasmids are currently being submitted to Addgene.

## Abbreviations

ACN: Acetonitrile
AGC: Automatic gain control
Ambic: Ammonium bicarbonate
bHLH: Basic helix-loop-helix
CTAD: C-terminal transactivation domain
DDA: Data dependent analysis
DTT: Dithiothreitol
FDR: False discovery rate
GO: Gene ontology
HCD: High energy collisional dissociation
HIF: Hypoxia inducible factor
HPLC: High pressure liquid chromatography
HTP: High through-put
IP: Immunoprecipitation
IPA: Ingenuity pathway analysis
LFQ: Label-free quantification
LTP: Low through-put
MS: Mass spectrometry
NCE: Normalised collision energry
NTAD: N-terminal transactivation domain
O_2_: Oxygen
ODDD: Oxygen dependent degradation domain
PAS-A/B: Per/Arnt/Sim domains A/B
PCR: Polymerase chain reaction
PD: Proteome discoverer
PSM: Peptide spectral match
PTM: Post translational modification
TFA: Trifluroacetic acid
TiO_2_: Titanium dioxide

## SUPPLEMENTAL FIGURES

**Supplemental Figure 1:**
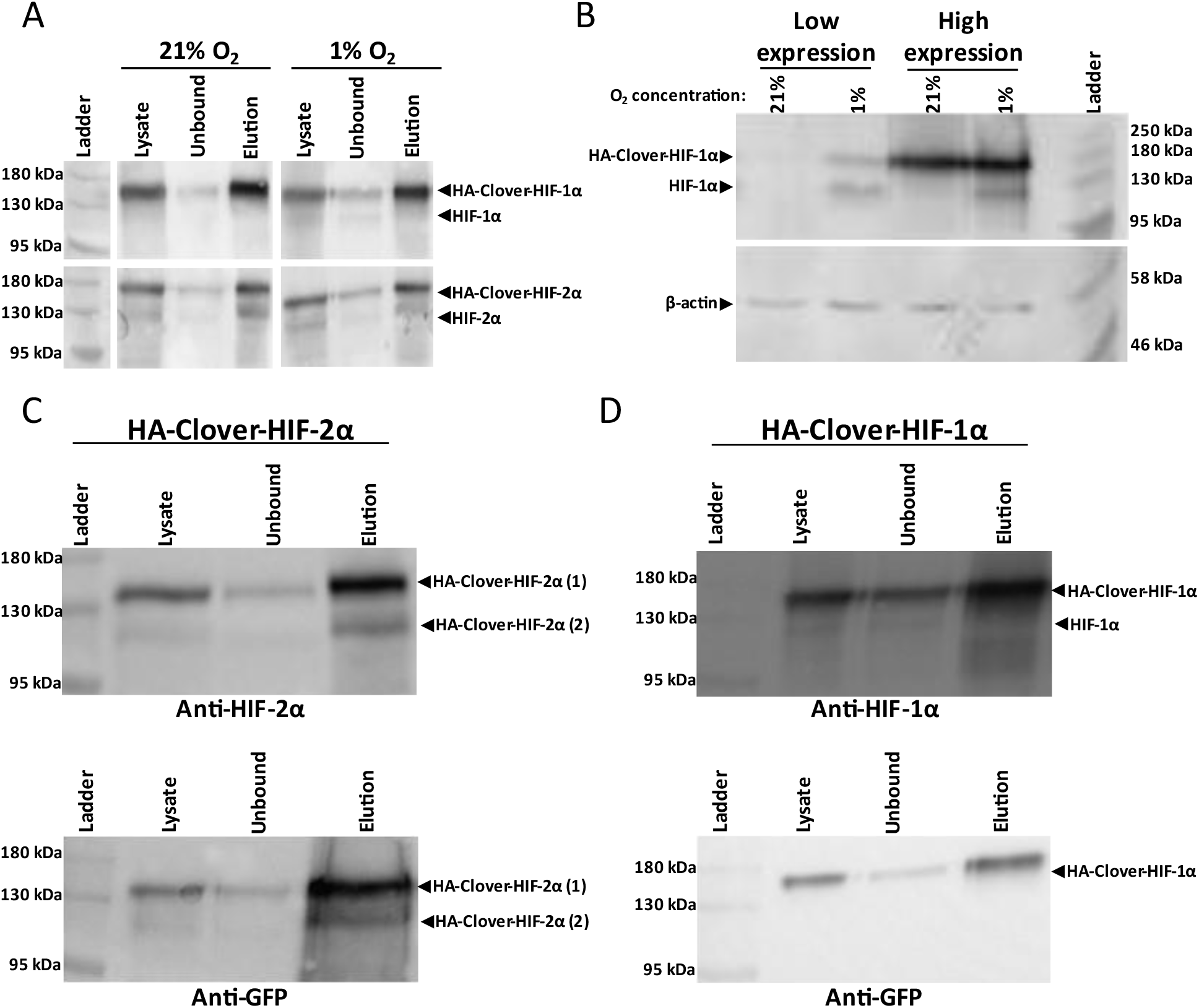
Immunoprecipitation of transfected HA-Clover-HIFα. (A) HeLa cells were transfected at the high expression level (1:1 ratio HA-Clover-HIFα:pcDNA(-)) and incubated at 21% or 1% O_2_ for 4 h prior to lysis. Western blot of pre-cleared lysate, unbound material and, post washing, RapiGest SF elution, for both HA-Clover-HIFα isoforms in 21% & 1% O_2_, probed with anti-HIF-1α or anti-HIF-2α. Equivalent loadings were analyzed for all samples. Representative of n=3, gaps indicate non-contiguous lanes. B) Comparison of high expression and low expression models. Low expression = 1:19 ratio HA-Clover-HIFα:pcDNA(-). Western blot probed with anti-HIF-1α and anti-β-actin. Equivalent loadings were analyzed for all samples. Representative of n=3. C) Identification that the higher and lower molecular weight bands observed are both HA-Clover-HIF-2α proteoforms. Western blots performed on the identical membrane with either Anti-HIF-2α or Anti-GFP antibodies. Cells transfected at the high expression and incubated at 1% O_2_ for 4 h prior to lysis. Equivalent loadings were analyzed for all samples. Representative of n=3. D) Identification that the higher molecular weight band observed is HA-Clover-HIF-1α and the lower molecular weight band observed is endogenous HIF-1α. Conditions as were in C.

**Supplemental Figure 2:**
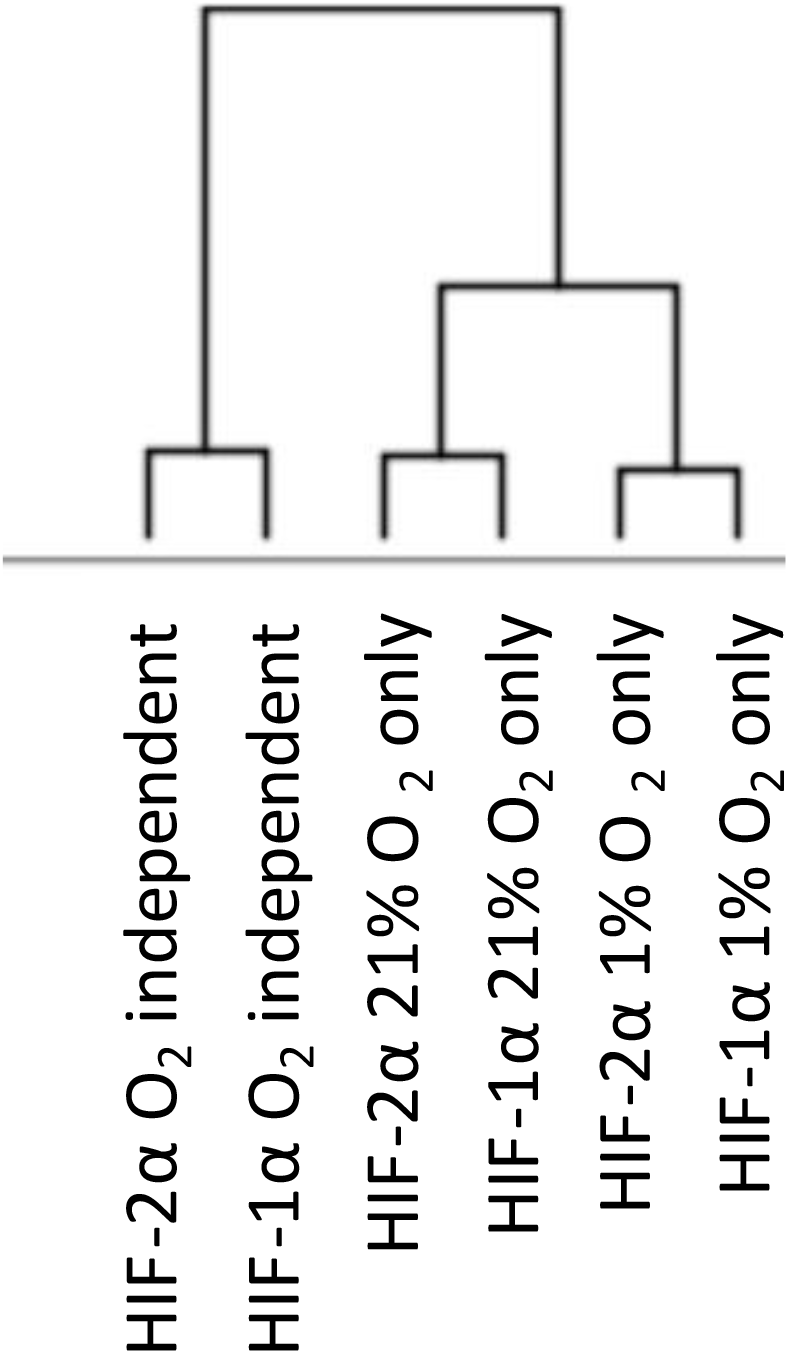
Hierarchal clustering of enriched pathways from HIF-1α and HIF-2α binding partners in different O_2_ conditions. IPA comparison analysis was performed on the binding partners as determined in Fig. 2A/3A.

**Supplemental Figure. 3:**
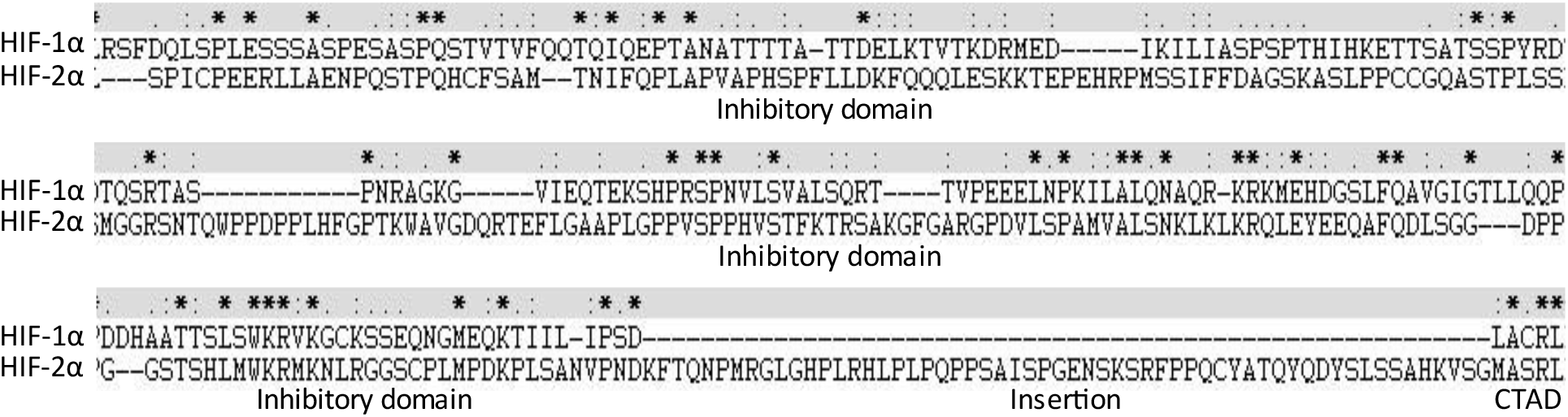
Sequence alignment of the inhibitory domains of HIF-1α versus HIF-2α (residues 576-785 and 543-821 respectively) using the MUSCLE multiple sequence alignment tool and viewed in Clustal X, * = identical residue: = similar residue properties. = less similar residue properties space = no similarity in residues (properties as determined by Clustal X) -= gap inserted to align sequences. Domains are labelled underneath the sequence alignments

**Supplemental Figure. 4:**
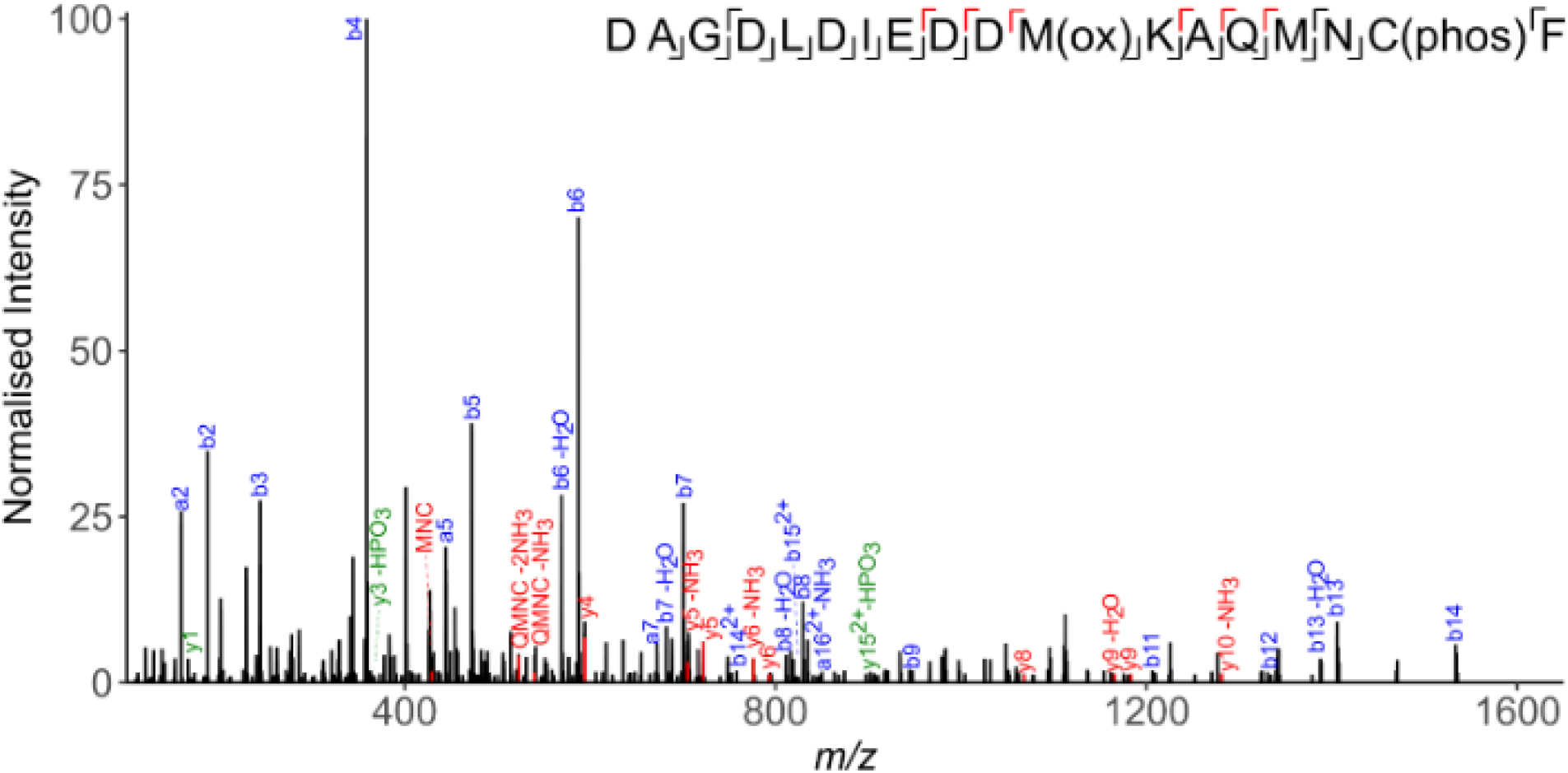
HIF-1α pCys90 annotated MS2 spectrum. MS/MS spectrum of the doubly charged peptide ion of m/z 1064.3971, fragmented using HCD. Peptide sequence is displayed with the annotated HCD product ions labelled, including the position of Met84 oxidation and Cys90 phosphorylation. a/b ions (blue), y ions (green) and site determining ions (red) are labelled. Only site determining internal fragment ions are displayed. Sequence and ions identified are displayed. Spectra representative of n=4.

